# Drug-induced change in transmitter identity is a shared mechanism generating cognitive deficits

**DOI:** 10.1101/2022.06.16.496480

**Authors:** Pratelli Marta, Anna M. Hakimi, Arth Thaker, Hui-quan Li, Swetha K. Godavarthi, Nicholas C. Spitzer

## Abstract

Cognitive deficits are a long-lasting consequence of drug use, yet the convergent mechanism by which classes of drugs with different pharmacological properties cause similar deficits is unclear. We find that both phencyclidine and methamphetamine, despite differing in their targets in the brain, impair memory by causing the same glutamatergic neurons in the medial prefrontal cortex to gain a GABAergic phenotype and decrease their expression of the vesicular glutamate transporter. Suppressing drug-induced gain of GABA with RNA-interference prevents the appearance of memory deficits. Drug-induced prefrontal hyperactivity drives this change in transmitter identity. Normalizing the activity of prefrontal glutamatergic neurons after drug-exposure reverses the gain of GABAergic phenotype and rescues the associated memory deficits. Increased activity of dopaminergic neurons in the ventral tegmental area is necessary and sufficient to produce the change in transmitter identity. The results reveal a shared and reversible mechanism by which exposure to different drugs causes cognitive deficits.

## Main

Brain impairments are often characterized by constellations of symptoms and behavioral alterations, some of which are shared across disorders. Cognitive deficits are found in mood and neuropsychiatric disorders such as drug-misuse, schizophrenia and depression, raising the possibility that shared mechanisms could produce these impairments under different conditions. To test this hypothesis, we investigated the effect of sub-chronic treatment with phencyclidine (PCP) and methamphetamine (METH), two drugs belonging to different classes of chemicals. PCP affects glutamatergic transmission by acting as an NMDA antagonist^1^, while METH affects signaling by dopamine and other monoamines^2^. Despite differing in their molecular targets in the brains and in some behavioral effects^1–7^, PCP and METH have been extensively studied for their ability to cause long-lasting cognitive deficits and mimic symptoms of schizophrenia^8–10^. While studies have focused on the actions of single drugs, little attention has been given to investigating the mechanisms of action that these drugs have in common. The process by which they generate similar behavioral impairments has been unknown. Understanding shared neuronal mechanisms underlying drug-induced cognitive deficits could foster development of effective treatments and be beneficial for a spectrum of disorders^11,12^.

When neuronal activity is altered for a sustained period, neurons can change the neurotransmitter they express, often switching from an excitatory to an inhibitory transmitter or vice versa and causing changes in behavior^13,14^. Linkage of drug-induced behavioral alterations to increased expression of specific transmitters in subcortical regions^15–17^ suggests that changes in transmitter identity could produce the behavioral effects of repeated exposure to drugs of abuse. Using a combination of genetic labeling strategies, RNA interference, chemogenetics and optogenetics, we investigated whether changes in cortical neuron transmitter phenotype are a shared mechanism underlying both PCP- and METH-induced cognitive deficits.

## Results

### Phencyclidine induces cognitive deficits by changing the transmitter phenotype of neurons in the medial prefrontal cortex

To determine whether cognitive deficits result from changes in neurotransmitter phenotype, we tested the effect of a 10-day treatment with PCP (10 mg/kg/day), which induces long-lasting cognitive impairments and recapitulates deficits observed in schizophrenia^10,18^. We focused on the medial prefrontal cortex (mPFC), which is a major hub for cognitive control^19,20^, and we examined the transmitter phenotype of glutamatergic neurons expressing vesicular glutamate transporter 1 (VGLUT1)^20^ because they represent the largest population in the mPFC. To identify these neurons following changes in transmitter profile, we labeled them permanently with a nuclear mCherry reporter using VGLUT1^CRE^::mCherry mice (Fig. 1a). In mice exposed to PCP, we identified 1198±59 mCherry^+^ neurons in the prelimbic subregion (PL) of the mPFC immunolabelled for GABA and 1096±81 labelled for its synthetic enzyme, glutamic acid decarboxylase 67 (GAD67) (Fig. 1b-d and Extended Data Fig. 1a-c). In control mice, there were only 622±45 and 643±22 of these neurons, indicating that PCP increased the number mCherry^+^/GABA^+^ and mCherry^+^/GAD67^+^ cells by 1.9- and 1.7-fold. PCP did not alter the number of mCherry-labeled PL neurons (Extended Data Fig. 1d), and no sign of apoptosis or neurogenesis was detected (Extended Data Fig. 2). These results suggested that PCP induces the synthesis and expression of GABA in PL glutamatergic neurons not previously expressing this transmitter.

**Fig. 1.**
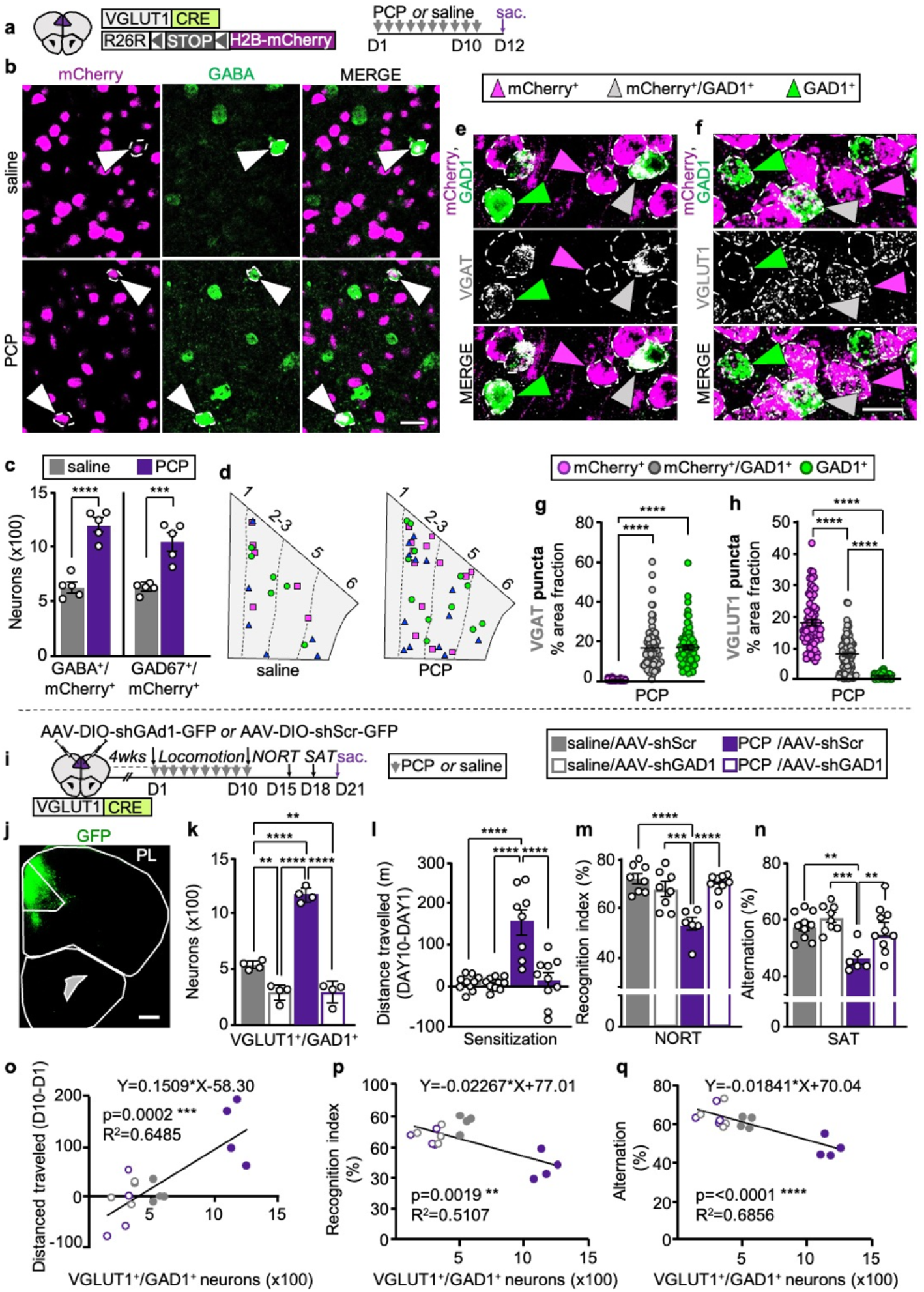
PCP changes the transmitter identity of PL glutamatergic neurons that in turn causes locomotor sensitization and cognitive deficits. **a**, Experimental protocol. **b**, PL neurons co-expressing mCherry and GABA (arrowheads). Scale bar, 25 μm. **c**, Quantification of mCherry^+^/GABA^+^ and mCherry^+^/GAD67^+^ neurons (*n*=5 mice). d, PL locations of glutamatergic neurons that co-expressed GABA or changed transmitter identity upon treatment with PCP. Cartoons were produced by superimposing the locations of neurons in three PL sections, each from a different mouse, all positioned at Bregma 1.94. Different shapes represent cells from different mice. **e,f**, Expression of VGAT and VGLUT1 mRNA puncta in PL neurons of PCP-treated mice. Scale bar, 20 μm. **g,h**, Quantification of VGAT and VGLUT1 expression measured as percent of cell area occupied by mRNA puncta across PL neuronal subtypes (*n*=25 cells/type/mouse for 3 mice). i, Experimental protocol. **j**, Expression of shGAD1-GFP in the PL. Scale bar, 500 μm. **k**, Quantification of neurons co-expressing VGLUT1 and GAD1 transcripts (*n*=4 mice). l-n, shGAD1 prevents PCP-induced deficits in locomotor sensitization (l, *n*=8-10 mice), in the NORT (m, *n*=6-9 mice) and SAT (n, *n*=6-10 mice). **o-q**, The number of PL VGLUT1^+^/GAD1^+^ neurons is positively correlated with locomotor sensitization and negatively correlated with performance in the NORT and SAT (*n*=4 mice). Statistical significance (**P<0.01, ***P<0.001, ****P<0.0001) was assessed using unpaired t-test (c), Kruskal-Wallis followed by Dunn’s test (**g,h**), two-way ANOVA followed by Tukey’s test (k-n), and linear regression and Pearson’s correlation analysis (**o-q**). Data are presented as mean ± SEM. See Supplementary Table 1 for detailed statistics.

We then examined the expression of the GABA vesicular transporter (VGAT) and VGLUT1 in mCherry^+^ neurons that gained GABA after PCP treatment or that co-expressed GABA in drug-naïve conditions. We used fluorescent *in situ* hybridization (FISH) to detect transcripts for mCherry, for the GABA synthetic enzyme (GAD1), and for either VGAT or VGLUT1. To reveal changes in expression levels, we selectively decreased the amplification for VGAT and VGLUT1 to obtain punctate staining (Fig. 1e,f). The density of neurons expressing both mCherry and GAD1 (mCherry^+^/GAD1^+^) was 1.6-fold higher in the PL of PCP-treated mice than in controls (6.6±0.5 neurons/mm^2^, saline; 10.6±0.9 neurons/mm^2^, PCP), mirroring the PCP-induced increase in the number of mCherry^+^/GAD67^+^ neurons (Fig. 1c) and suggesting that PCP induced gain of GAD1 in mCherry^+^ neurons that were not expressing it earlier. mCherry^+^/GAD1^+^ neurons in the PL of PCP-treated mice expressed VGAT at the level of GABAergic neurons (labeled with GAD1 and not mCherry) (Fig. 1g). At the same time, the expression level of VGLUT1 in mCherry^+^/GAD1^+^ neurons decreased by ∼55% compared to that of glutamatergic cells expressing mCherry and not GAD1 in PCP-treated mice (Fig. 1h). Neurons co-expressing mCherry and GABA in drug-naïve conditions also showed high expression levels of VGAT and low VGLUT1 (46% less VGLUT1 than in purely glutamatergic neurons expressing only mCherry), as evident from analyses of these cells in saline-treated controls (Extended Data Fig. 1e,f). Thus, both the glutamatergic neurons that gained GABA after PCP-exposure, as well as those expressing GABA in drug-naïve conditions express high levels of VGAT and lowered levels of VGLUT1.

We next asked whether PCP-treatment affects the transmitter phenotype of PL GABAergic neurons. No difference was observed in the number of neurons expressing GABA and not mCherry between PCP-treated animals and controls (8594±340 vs 8837±271) (Extended Data Fig. 1g). However, PCP and other NMDA receptor antagonists have been shown to reduce the expression of GAD67 and parvalbumin in prefrontal cortex parvalbumin-positive (PV^+^) interneurons^21,22^. Because variability in the number of GABAergic neurons scored could have prevented the detection of loss of GABA from a small number of PV^+^ neurons, we quantified the number of PV^+^ neurons expressing GAD67 after PCP-treatment by permanently labeling them with a PV^CRE^::TdTomato mouse line. PCP caused 223±45 TdTomato^+^ neurons (6% of the TdTomato^+^ population) to stop expressing GAD67 (Extended Data Fig. 1h-k), indicating that PCP treatment reduces the expression of GABAergic markers in a subpopulation of PV^+^ neurons.

To investigate whether glutamatergic neurons that have gained GABA contribute to cognitive deficits, we selectively suppressed GABA expression in PL glutamatergic neurons by injecting a Cre-dependent adeno-associated virus (AAV) expressing shRNA for GAD1 (AAV-DIO-shGAD1-GFP or AAV-DIO-shScr-GFP as control) in the PL of VGLUT1^CRE^ mice before exposure to PCP (Fig. 1i,j). shGAD1 suppressed GABA expression in transfected cells (Extended Data Fig. 3a-c), and reduced the number of PL neurons co-expressing GAD1 and VGLUT1 transcripts in both PCP- and saline-treated mice to half of that in saline-ShScr controls (Fig. 1k). Having efficiently overridden PCP-induced gain of GABA, we examined the impact of shGAD1 on PCP-induced behavior. While shGAD1 did not affect PCP-induced hyperlocomotion on the first day of treatment, it prevented appearance of locomotor sensitization after a 10-day PCP-exposure (Fig. 1l and Extended Data Fig. 3d), indicating that PCP-induced gain of GABA is required for sensitization to the acute locomotor effect of the drug. We next focused on deficits in recognition and working memory, since these behaviors are affected by repeated exposure to PCP^18,23^ and are regulated by the PL^24,25^. shGAD1 prevented PCP-induced impairments of recognition memory in the novel object recognition test (NORT) (Fig. 1m) and deficits in spatial working memory in the spontaneous alternation task (SAT) (Fig. 1n). Neither PCP-treatment nor shGAD1 changed exploratory behaviors in the NORT and in the SAT (Extended Data Fig. 3e,f). shGAD1 did not affect the behavioral performances of saline-treated controls (Fig. 1l-n), suggesting that glutamatergic neurons gaining GABA upon PCP-exposure, but not those co-expressing GABA before drug-exposure, mediated drug-induced locomotor sensitization and memory impairments. The number of PL VGLUT1^+^/GAD1^+^ neurons was positively correlated with locomotor sensitization, and negatively correlated with object recognition and working memory performance (Fig. 1o-q). Overall, these data indicate that PCP-induced gain of GABA in PL glutamatergic neurons produces these behavioral alterations.

### Methamphetamine changes the transmitter phenotype of the same prelimbic neurons affected by phencyclidine

Because METH causes memory deficits similar to those induced by PCP^25–27^, we asked whether METH-treatment would also affect the transmitter identity of PL glutamatergic neurons. Resembling the effect of PCP, 10 days of METH-treatment (1 mg/kg/day) increased the number of mCherry^+^ PL neurons co-expressing GABA and GAD67 by 1.7- and 1.9-fold (Fig. 2a-d and Extended Data Fig. 4a), without changing the number of mCherry^+^ and GABA^+^/mCherry^-^ cells (Extended Data Fig. 4b) or inducing apoptosis or neurogenesis (Extended Data Fig. 2). We next used FISH to examine the expression of the VGAT and VGLUT1 in mCherry^+^ neurons that co-expressed or gained GABA. Similar to the effects of PCP-treatment, mCherry^+^/GAD1^+^ neurons in the PL of METH-treated mice expressed a high level of VGAT (equal to that of neurons expressing GAD1 but not mCherry) (Fig. 2e). The expression level of VGLUT1 decreased by ∼76% compared to that of glutamatergic cells expressing mCherry and not GAD1 (Fig. 2f). The number of mCherry^+^ neurons gaining expression of GABA, GAD67 and VGAT and reducing their expression level of VGLUT1 after METH-treatment resembled the number observed in PCP-treated mice, indicating that both drugs affect PL glutamatergic neurons similarly.

**Fig. 2.**
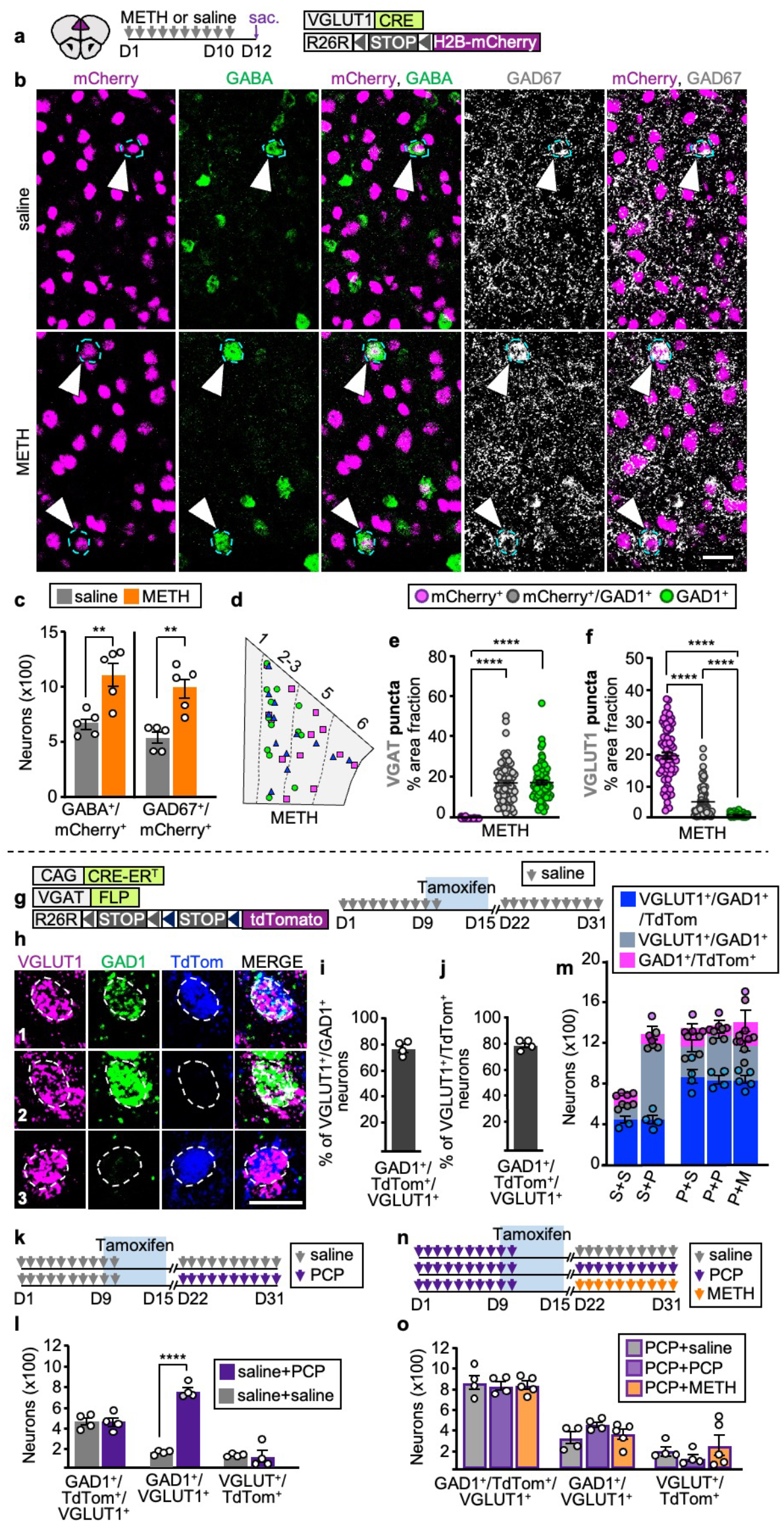
METH causes the same change in PL glutamatergic neuron transmitter phenotype. **a**, Experimental protocol. **b**, PL neurons co-expressing mCherry, GABA, and GAD67 (arrowheads). Scale bar, 25 μm. **c**, Quantification of PL neurons co-expressing mCherry and GABA or GAD67 (*n*=5 mice). **d**, PL locations of glutamatergic neurons that co-expressed GABA or changed transmitter identity upon treatment with METH generated as in Fig. 1d. **e**,**f**, VGAT and VGLUT1 expression measured as percent of cell area occupied by mRNA puncta (*n*=25 cells/type/mouse for 3 mice). **g**, Mouse line used for tamoxifen-inducible genetic labeling and experimental protocol to test efficiency and specificity. **h**, Examples of neurons that are VGLUT1^+^/GAD1^+^/TdTom^+^ (1), VGLUT1^+^/GAD1^+^ (2), and VGLUT1^+^/TdTom^+^ (3). Scale bar, 20 μm. **i**,**j**, Efficiency (77%, 1788 VGLUT1^+^/GAD1^+^/TdTom^+^ / 2328 VGLUT1^+^/GAD1^+^ neuroåns) and specificity (79%, 1788 VGLUT1^+^/GAD1^+^/TdTom^+^ / 2268 VGLUT1^+^/TdTom^+^ neurons) of this approach in labelling VGLUT1^+^/GAD1^+^ neurons with the TdTomato line (*n*=4 mice). These data were obtained from the same animals shown in (**l**). **k**, Experimental protocol to distinguish VGLUT1 neurons co-expressing GAD1 in drug-naïve conditions from those gaining GAD1 upon PCP treatment. **l**, Quantification of the neurons shown in (**h**) in mice treated as described in (**k**) (*n*=4 mice). **m**, Cumulative number of neurons that are VGLUT1^+^/GAD1^+^/TdTom^+^, VGLUT1^+^/GAD1^+^, and VGLUT1^+^/TdTom^+^ from (**l**) and (**o**) (S+S: saline+saline; S+P: saline+PCP; P+S: PCP+saline; P+P: PCP+PCP; P+M: PCP+METH). **n**, Experimental protocol to learn whether serial administration of PCP and METH changes the transmitter phenotype of the same number of neurons as PCP alone, causes neurons that have gained GAD1 to lose it, or enables other neurons to gain GAD1. **o**, Quantification of neurons shown in (**h**) in mice treated as described in (**n**) (*n*=4-5 mice). Statistical significance (**P<0.01, ****P<0.0001) was assessed using unpaired t-test and Mann Whitney U (**c**),Kruskal-Wallis followed by Dunn’s test (**e**,**f**), and two-way ANOVA followed by Tukey’s test (**l**,**o**). Data are presented as mean ± SEM. See Supplementary Table 1 for detailed statistics.

Glutamatergic neurons that co-express GABA or gain it after treatment with either drug were most prevalent in layer 2/3 and layer 5 of the PL (Extended Data Fig. 4c). These PL layers innervate the nucleus accumbens (NAc)^28^, which modulates behaviors that are affected by repeated intake of PCP or METH^29–31^. To determine whether neurons that switch transmitter identity project to the NAc, we injected fluoro-gold (FG) into the NAc of VGLUT1^CRE^::mCherry mice, treated them with PCP, METH or saline, and screened the PL for mCherry^+^/GABA^+^ neurons expressing the retrograde tracer (Extended Data Fig. 4d-f). In both PCP- and METH-treated mice, ∼0.9% of FG^+^ neurons were mCherry^+^/GABA^+^. Such cells were less frequent in controls (∼0.3% of the total number of FG^+^ neurons) (Extended Data Fig. 4g), indicating that neurons changing transmitter identity with drug-treatment project to the NAc.

Since both PCP and METH affect the transmitter phenotype of PL glutamatergic neurons that have the NAc as a shared downstream target, we asked whether both drugs change the transmitter identity of the same cells. If PCP and METH changed the transmitter identity of different cells, administering the two drugs one after the other should induce gain of GABA in neurons that have not gained it after treatment with the first drug. To determine whether this was the case, we genetically labeled neurons expressing GABAergic markers during the interval between the delivery of PCP and METH, using VGAT^FLP^::CreER^T^::TdTomato^cON/fON^ mice (see Methods) in which neurons expressing VGAT at the time of tamoxifen administration are permanently labeled with TdTomato (Fig. 2g). We first injected tamoxifen in saline-treated controls and determined that TdTomato labels neurons co-expressing VGLUT1 and GAD1 in drug-naïve conditions with 77% efficiency and 79% specificity (Fig. 2h-j).

We then used this labelling approach to distinguish neurons expressing GAD1 in drug-naïve mice from those gaining it upon drug-exposure, by administering mice with PCP after saline- and tamoxifen-treatment (Fig. 2k). PCP administration increased the total number of PL VGLUT1^+^/GAD1^+^ neurons 2-fold compared to controls (1188±23, saline+PCP; 582±27, saline+saline) (Fig. 2l), in line with previous findings (Fig. 1k). We detected no differences in the number of VGLUT1^+^/GAD1^+^/TdTomato^+^ neurons (441±43, saline+PCP; 447±34, saline+saline) and VGLUT1^+^/TdTomato^+^ neurons (99±61, saline+PCP; 120±8, saline+saline) between saline+PCP mice and saline+saline controls (Fig. 2l,m). Changes in these numbers would have indicated that drug-treatment caused some glutamatergic neurons co-expressing GAD1 in drug-naïve conditions to lose expression of GAD1. The results indicate that PCP induces expression of GAD1 in a population of PL neurons that were not previously expressing it, without affecting the transmitter phenotype of cells co-expressing GAD1 and VGLUT1 in drug-naïve conditions.

We next used VGAT^FLP^::CreER^T^::TdTomato^cON/fON^ mice to determine if PCP and METH cause the same neurons to switch transmitter phenotype. Mice were treated first with PCP followed by tamoxifen administration, and then treated with either saline, PCP or METH (Fig. 2n). Across treatment groups the total number VGLUT1^+^/GAD1^+^ neurons was unchanged (1169±46, PCP+saline; 1273±69, PCP+PCP; 1177±45, PCP+METH) (Fig. 2m,o), indicating that consecutive administration of drugs does not cause additional glutamatergic neurons to gain GAD1. Furthermore, we did not detect differences in the number of VGLUT1^+^/GAD1^+^/TdTomato^+^ neurons (861±73, PCP+saline; 831±43, PCP+PCP; 832±38, PCP+METH) nor in the number of VGLUT1^+^/TdTomato^+^ neurons (180±44, PCP+saline; 108±41, PCP+PCP; 237±103, PCP+METH) (Fig. 2m,o). A decrease in the first population and an increase in the second population would have indicated loss of GAD1 from some VGLUT1^+^ neurons and gain of GAD1 in another population of VGLUT1^+^ neurons. These results indicated that consecutive administration of PCP and METH does not cause gain of GAD1 by additional neurons, nor induces neurons that gained GAD1 upon PCP-treatment to revert to their original transmitter phenotype. Thus, PCP and METH change the transmitter identity of a largely overlapping population of PL neurons.

### Drug-induced prelimbic hyperactivity mediates the change in transmitter phenotype and linked cognitive deficits

Demonstration that both PCP and METH have the same effect on the transmitter phenotype of the same PL glutamatergic neurons prompted investigation of the underlying mechanism of drug action. Increased neuronal activity can cause neurons to change the transmitter they express^13,32,33^. Could PCP and METH induce alterations in PL activity that mediate the switch in PL neuron transmitter phenotype? PCP and METH increased c-fos expression in PL glutamatergic neurons by 3.8- and 3.5-fold after a single injection and by 2.6- and 3.7-fold throughout a 10-day treatment (Extended Data Fig. 5a-d,f,g). This PL hyperactivity was still present 2 days after the end of drug-treatment (Extended Data Fig. 5i,j). To determine whether this increase in activity promoted the switch in transmitter phenotype, we tested whether suppression of PL hyperactivity would prevent glutamatergic neurons from gaining GABA. Glutamatergic cells in the PL receive perisomatic inhibition from local PV^+^ interneurons, which do not show changes in c-fos expression after administration of PCP or METH (Extended Data Fig. 5e,h,k). We hypothesized that chemogenetic activation of PV^+^ neurons would suppress drug-induced hyperactivity of glutamatergic cells^34,35^. To test this idea, we expressed the chemogenetic receptor PSAML-5HT3HC in mPFC PV^+^ neurons and administered PSEM^308^ immediately before acute injection of either PCP or METH (Extended Data Fig. 6a-e). While GFP^+^ neurons infected with the virus showed high c-fos expression after PSEM^308^ treatment, consistent with their expected activation (Extended Data Fig. 6f), the PCP- or METH-induced increase in c-fos expression in PL glutamatergic neurons was suppressed (Extended Data Fig. 6g-i).

We then combined chemogenetic activation of PL PV^+^ interneurons with either PCP- or METH-administration for the duration of drug-treatment (Fig. 3a,b). The number of VGLUT1^+^/GAD1^+^ neurons in the PL of PCP- and METH-treated mice that received PSEM^308^ was half of that of mice that did not (586±18 and 611±23 vs 1262±66 and 1222±12) and was indistinguishable from that of saline-treated controls (Fig. 3c,d). Thus, suppression of drug-induced PL hyperactivity is sufficient to prevent glutamatergic neurons from switching their transmitter identity, indicating that hyperactivity mediates the change in transmitter phenotype.

**Fig. 3.**
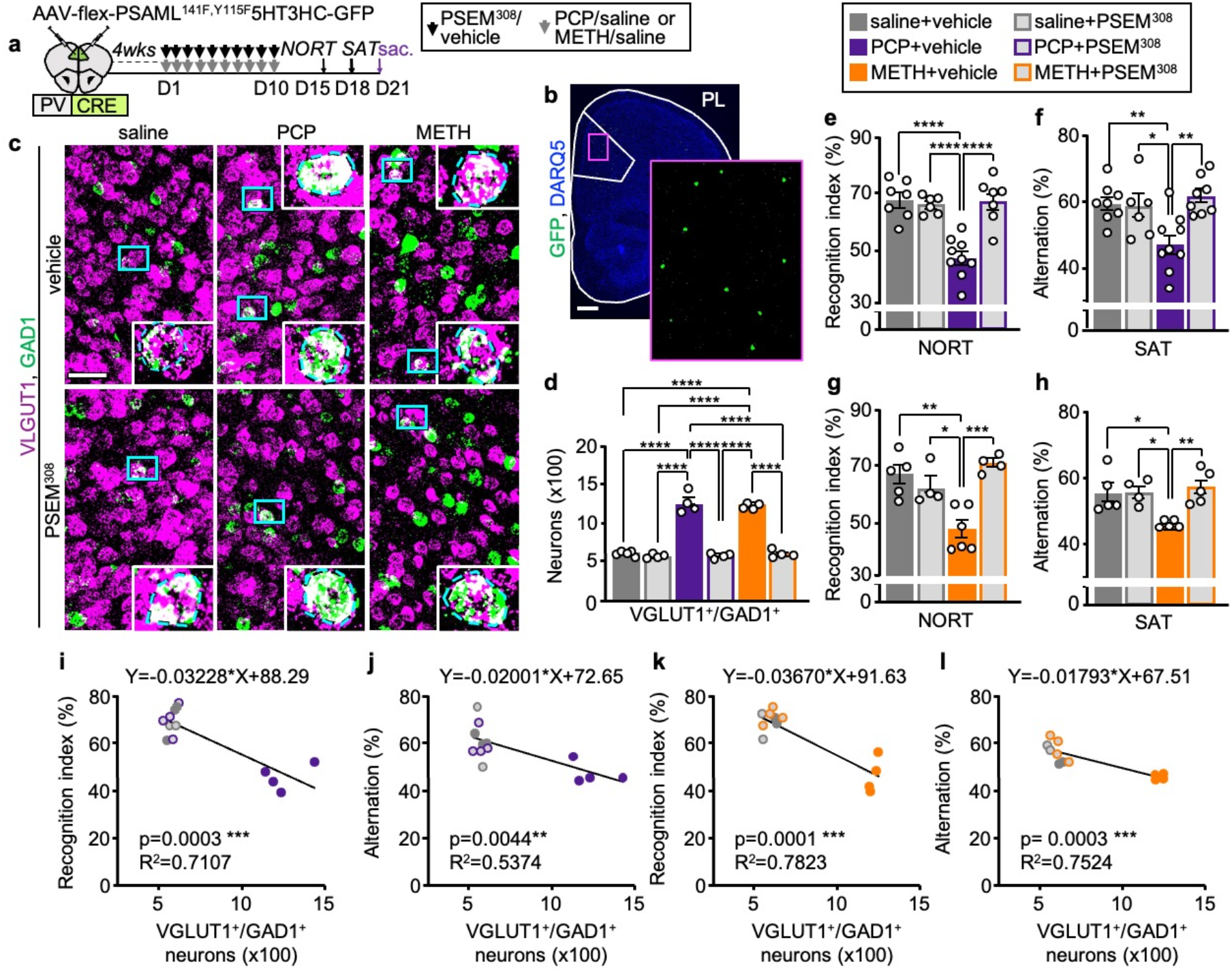
Suppression of drug-induced PL hyperactivity prevents the change in transmitter identity and drug-induced behavioral alterations. **a**, Experimental protocol. **b**, Expression of PSAML-5HT3HC-GFP in the PL. Scale bar, 500 μm. **c**, VGLUT1 and GAD1 expression in the PL following saline/drug treatment combined with chemogenetic activation of PV^+^ neurons. (Blue rectangles) neurons co-expressing VGLUT1 and GAD1 illustrated at high magnification in the insets. Scale bar, 50 μm. **d**, Quantification of **c** (*n*=4-5 mice). **e**-**h**, Chemogenetic activation of PV^+^ neurons during drug treatment prevents the deficits in the NORT and SAT induced by both PCP and METH (*n*=4-9 mice). **i**-**l**, The number of PL VGLUT1+/GAD1+ neurons correlates with performance in the NORT and SAT (*n*=2-4 mice). Statistical significance (*P<0.05, **P<0.01, ***P<0.001, ****P<0.0001) was assessed using two-way ANOVA followed by Tukey’s test (**d**-**h**) and linear regression and Pearson’s correlation analysis (**i**-**l**). Data are presented as mean ± SEM. See Supplementary Table 1 for detailed statistics.

We now tested whether blocking the change in transmitter phenotype through chemogenetic activation of PV^+^ neurons was sufficient to prevent drug-induced cognitive deficits. In mice treated with PSEM^308^, drug-induced hyperlocomotion was absent on both the first and last days of treatment (distance traveled on DAY1: 30±16m, saline+vehicle; 29±18m, saline+PSEM^308^; 173±69m, PCP+vehicle; 45±25m, PCP+PSEM^308^; 137±37m, METH-saline; 35±14m, METH+PSEM^308^. Distance traveled on DAY10: 32±19m, saline+vehicle; 34±30m, saline+PSEM^308^; 309±67m, PCP+vehicle; 67±26m, PCP+PSEM^308^; 192±30m, METH-saline; 57±18m, METH+PSEM^308^). These results are consistent with suppression of acute drug-induced hyperactivity of PL glutamatergic neurons ^36^.

Suppressing PL activity prevented both PCP- and METH-induced appearance of memory deficits in both the NORT and the SAT (Fig. 3e-h), without influencing exploratory behaviors (Extended Data Fig. 6j-n). The number of VGLUT1^+^/GAD1^+^ neurons in the PL was negatively correlated with the performance in the NORT and SAT (Fig. 3i-l). These results suggest that chemogenetic activation of PV^+^ neurons specifically affects the performance of drug-treated mice by preventing the change in the transmitter phenotype of glutamatergic neurons in the PL.

### Normalizing prelimbic neuron activity after drug-exposure reverses the change in transmitter identity and the associated behavioral alterations

PL glutamatergic neurons that change their transmitter upon exposure to PCP or METH retain the GABAergic phenotype for at least 11 days of drug washout (Fig. 3d, and PCP+saline group in Fig. 2o). As PCP- and METH-induced memory deficits are also long-lasting^18,37^, we asked whether the persistence of behavioral deficits is linked to retention of the GABAergic phenotype and whether both can be reversed. Clozapine, a powerful antipsychotic drug, reverses PCP-induced deficits in the NORT^18^, leading us to investigate whether it also reverses the change in transmitter phenotype. VGLUT1^CRE^::mCherry mice that received PCP displayed 1.94 fold more mCherry^+^/GABA^+^ PL neurons than controls, 17 days after the end of PCP-treatment, indicating that neurons had maintained the newly acquired GABAergic phenotype (Fig. 4a-d). In mice that received clozapine treatment after PCP, the number of mCherry^+^/GABA^+^ neurons was reduced compared to that of mice treated with PCP alone (559±55 vs 1124±94) and was not different from that of saline-treated controls (Fig. 4a-c). Clozapine did not affect the number of mCherry^+^/GABA^+^ neurons in saline-treated mice, suggesting that this drug selectively reverses the PCP-induced change in glutamatergic neuron transmitter identity. We found that clozapine rescued PCP-induced memory deficits in the NORT and SAT, without affecting the behavioral performance of controls (Fig. 4d-g and Extended Data Fig.7 a-d).

**Fig. 4.**
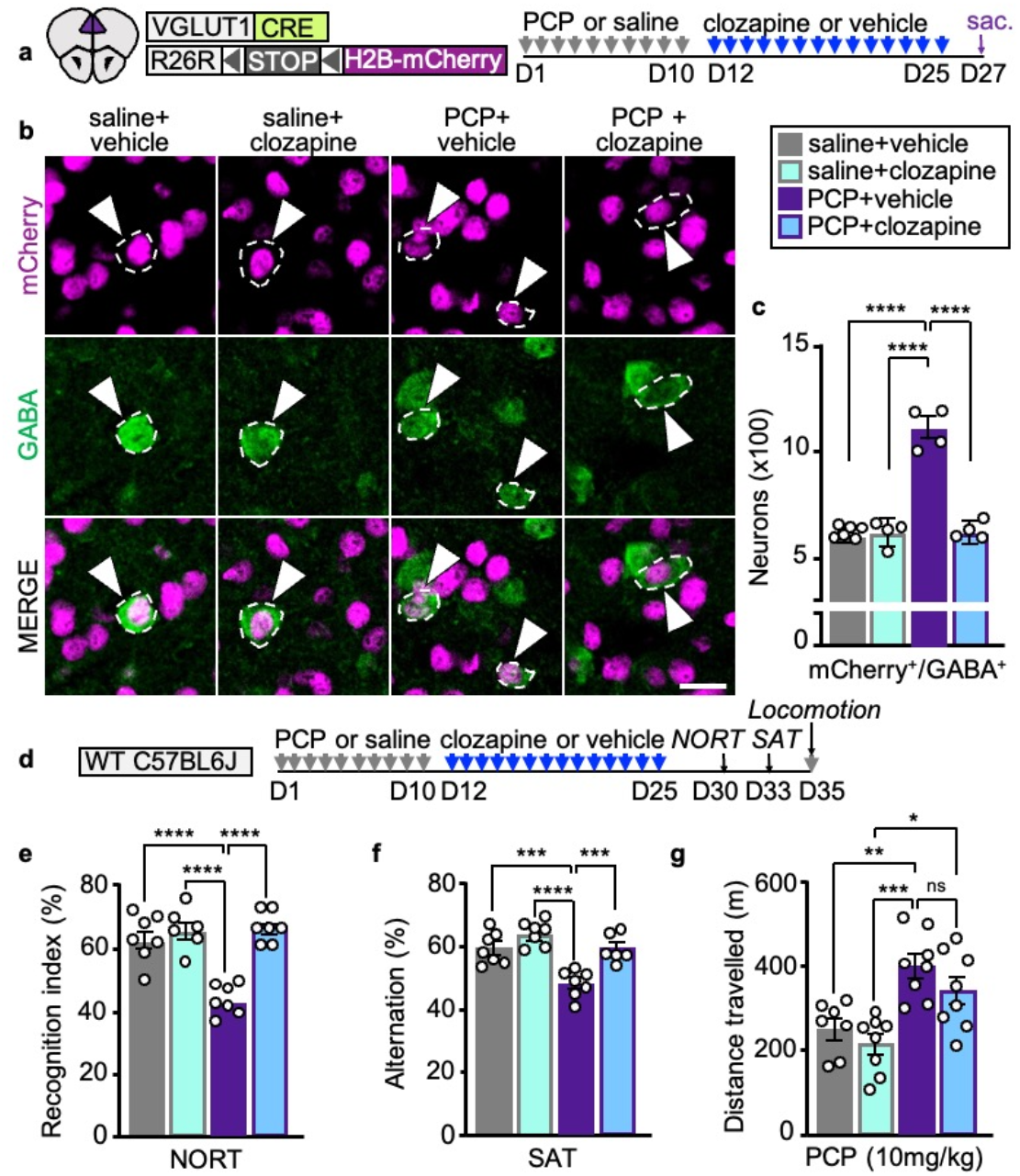
Clozapine treatment reverses PCP-induced gain of GABA in PL neurons and associated behaviors. **a**, Experimental protocol. **b**, PL neurons co-expressing mCherry and GABA (arrowheads). Scale bar 20 μm. **c**, Quantification of PL neurons co-expressing mCherry and GABA after treatment with PCP or saline followed by clozapine or vehicle (*n*=4-6 mice). **d**, Experimental protocol. **e**-**g**, Clozapine reverses PCP-induced deficits in the NORT and SAT but not locomotor sensitization after a single PCP challenge (*n*=6-8 mice). Statistical significance (*P<0.05, **P<0.01, ***P<0.001, ****P<0.0001) was assessed using two-way ANOVA with Tukey’s multiple-comparisons test (**c**,**e**-**g**). Data are presented as mean ± SEM. See Supplementary Table 1 for detailed statistics.

We next investigated the mechanisms underlying clozapine-induced reversal of the switch in transmitter identity. Because clozapine suppresses the acute PCP-induced increase in PL c-fos expression^38,39^, we asked whether reversal of the gain of GABA depends on suppression of neuronal activity. After drug-treatment, the number of c-fos^+^ glutamatergic neurons was 2.1- and 2.8-fold higher in PCP- and METH-treated mice compared to controls (Extended Data Fig. 5i-k). Administration of clozapine after PCP-treatment returned c-fos expression to baseline (Extended Data Fig. 7e-h), suggesting that PL hyperactivity during drug-washout is necessary to maintain the newly acquired transmitter phenotype. If this were the case, suppressing PL hyperactivity after the transmitter switch has occurred could be expected to reverse the change. To test this hypothesis, we chemogenetically activated PV^+^ neurons for 10 days to normalize c-fos expression in the PL of PCP- or METH-treated mice after the change in transmitter phenotype had taken place (Extended Data Fig. 8a-d). More than 3 weeks after the end of drug-treatment, glutamatergic neurons in the PL of both PCP and METH-treated mice still displayed the drug-induced GABAergic phenotype (Fig. 5a-c). Normalizing PL activity decreased the number of VGLUT1^+^/GAD1^+^ neurons in the PL of PCP and METH-treated mice to the level of controls (588±27 and 575±9 vs 1225±38 and 1071±42) (Fig. 5a-c). Thus, PL neuronal activity maintains the transmitter switch once it has been induced. Chemogenetically activating PL PV^+^ interneurons after the change in transmitter phenotype had occurred also rescued memory deficits in the NORT and SAT and suppressed locomotor sensitization to both PCP and METH (Fig. 5d-i and Extended Data Fig. 8e-k). Overall, these data show that suppressing PL hyperactivity following drug-exposure reverses the change in transmitter phenotype and the associated behavioral alterations.

**Fig. 5.**
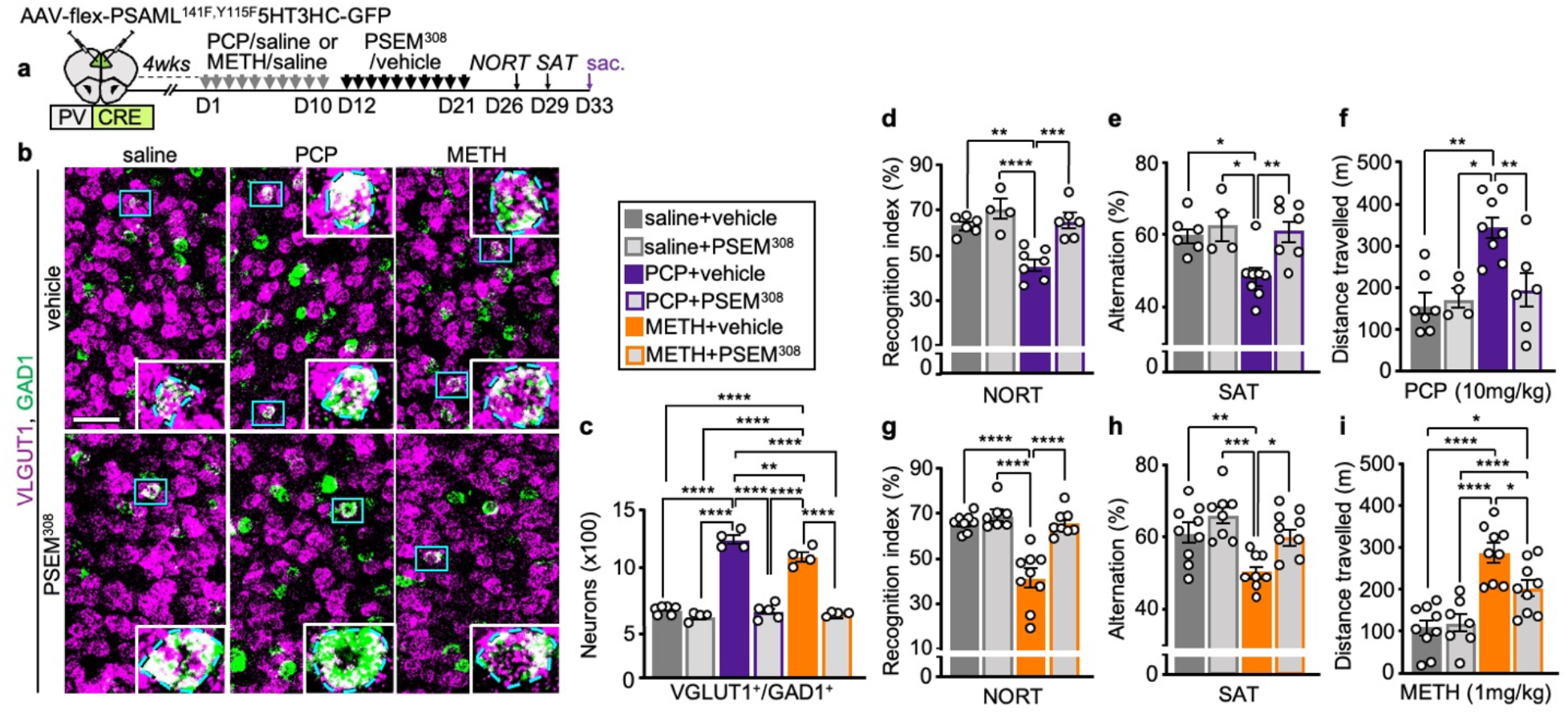
Suppression of drug-induced PL hyperactivity after the end of drug-treatment reverses the change in transmitter identity and drug-induced behavioral alterations. **a**, Experimental protocol. **b**, PL expression of VGLUT1 and GAD1 after exposure to saline/drug treatment followed by chemogenetic activation of PV^+^ neurons. (Blue rectangles) higher magnification of VGLUT1^+^/GAD1^+^ neurons in insets. Scale bar, 50 μm. **c**, Quantification of (**b**) (*n*=4-6 mice). **d**-**i**, PCP- and METH-induced locomotor sensitization and deficits in the NORT and SAT are reversed by sustained activation of PV^+^ neurons (*n*=4-9 mice). Statistical significance (*P<0.05, **P<0.01, ***P<0.001, ****P<0.0001) was assessed using two-way ANOVA followed by Tukey’s test (**c**-**i**). Data are presented as mean ± SEM. See Supplementary Table 1 for detailed statistics.

### Signaling by dopaminergic neurons is necessary and sufficient to change the transmitter identity of prelimbic neurons

PCP, METH, and other addictive substances increase phasic firing of dopaminergic neurons in the ventral tegmental area (VTA)^40,41^ and increase the levels of extracellular dopamine (DA) in the striatum and prefrontal cortex^42,43^. Could signaling by dopaminergic neurons in the VTA be a common mediator of the PCP- and METH-induced transmitter switch? To address this question, we tested whether suppressing the activity of VTA dopaminergic neurons during treatment with PCP or METH affects the number of PL neurons that switch transmitter phenotype. We expressed the PSAML-GlyR chemogenetic receptor in the VTA of DAT^CRE^ mice (Extended Data Fig. 9a-d). Administration of PSEM^308^ before drug-injection suppressed the acute PCP- and METH-induced increase in c-fos^+^ dopaminergic neurons (Extended Data Fig. 9e,f). Combining VTA suppression with drug administration for the entire duration of treatment, by co-administration of PSEM^308^ and PCP or METH, prevented the increase in the number of PL VGLUT1^+^/GAD1^+^ neurons (Fig. 6a-d). These results show that drug-induced increase in activity of VTA dopaminergic neurons is required for PL neurons to change their transmitter phenotype upon treatment with PCP or METH.

**Fig. 6:**
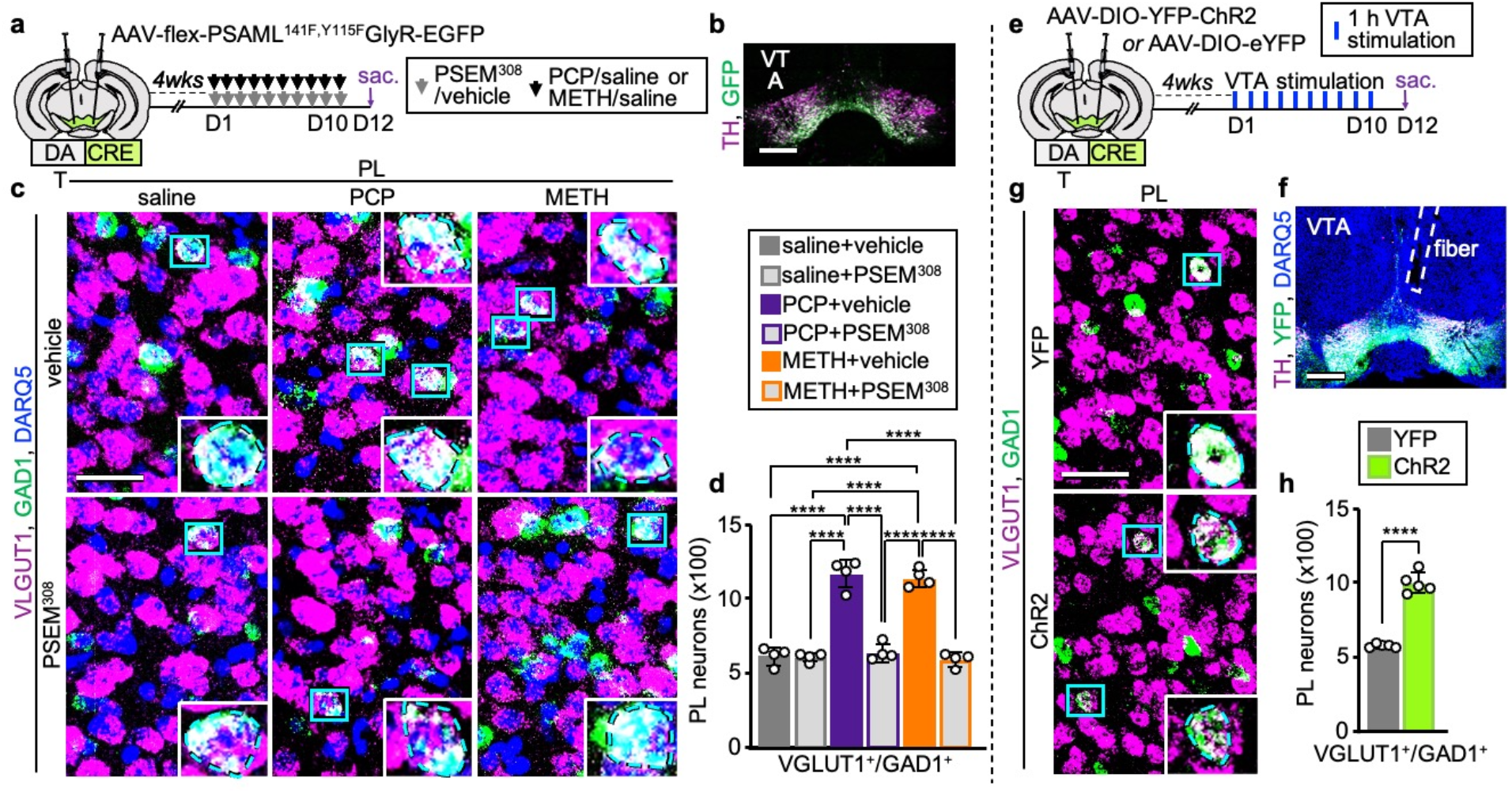
Activity of VTA DA neurons is necessary and sufficient to induce PL glutamatergic neurons to switch their transmitter identity. **a**, Experimental protocol. **b**, Expression of PSAML-GlyR-GFP in the VTA. Scale bar, 500 μm. **c**, VGLUT1 and GAD1 expression in the PL following saline/drug treatment combined with chemogenetic inhibition of VTA dopaminergic neurons. (Blue rectangles) neurons co-expressing VGLUT1 and GAD1 illustrated at high magnification in the insets. Scale bar, 50 μm. **d**, Quantification of VGLUT1^+^/GAD1^+^ neurons in the PL (*n*=4 mice). **e**, Experimental protocol. **f**, Expression of ChR2-YFP in the VTA and fiber location. Scale bar, 500 μm. **g**, VGLUT1 and GAD1 expression in the PL following 10 days of VTA optogenetic stimulation. (Insets) Higher magnification of neurons co-expressing VGLUT1 and GAD1. Scale bar, 50 μm. **h**, Quantification of VGLUT1^+^/GAD1^+^ neurons in the PL (*n*=5 mice). Scale bar, 50 μm. Statistical significance (****P<0.0001) was assessed using two-way ANOVA with Tukey’s multiple-comparisons test (**d**) and unpaired t-test (**h**). Data are presented as mean ± SEM. See Supplementary Table 1 for detailed statistics.

It remained unclear, however, whether stimulation of dopaminergic neurons in the VTA is by itself sufficient to induce PL neurons to switch transmitter phenotype, or whether other effects of PCP or METH are involved. Phasic firing of VTA neurons can be mimicked by optogenetic stimulation of VTA dopaminergic neurons^44^. We asked whether repeated optogenetic stimulation of VTA dopaminergic neurons is sufficient to induce PL glutamatergic neurons to switch transmitter identity. We expressed ChR2-YFP (or YFP as control) in VTA DAT^CRE^ neurons and implanted an optic fiber above the VTA (Fig. 6e,f). Administration of 80 sets of laser stimulation (each consisting of 30 bursts of five 8.5 ± 1 mW pulses of 4 ms duration at 20 Hz) over the course of 1h increased the number of VTA dopaminergic neurons expressing c-fos by 7.3-fold in mice expressing ChR2 (Extended Data Fig. 10a-d). Expression of c-fos in PL glutamatergic neurons was also increased by 2.7-fold, resembling the effect of a single dose of PCP or METH (Extended Data Fig. 10e-g). We then exposed mice to 1h of VTA stimulation per day for 10 days and analyzed the transmitter phenotype of PL glutamatergic neurons (Fig. 6e). Remarkably, the number of VGLUT1^+^/GAD1^+^ was 1.7-fold higher in ChR2-expressing mice compared to controls (Fig. 6g,h and Extended Data Fig. 10h-j), demonstrating that phasic firing of dopaminergic neurons in the VTA changes the transmitter phenotype of PL glutamatergic neurons. These findings establish signaling by dopaminergic neurons in the VTA as a common mediator for PCP- and METH-induced gain of GABA in PL glutamatergic neurons and suggest that exposure to other addictive substances that activate the VTA could produce similar effects.

## Discussion

We show that a change in the transmitter identity of PL glutamatergic neurons is a shared mechanism underlying both PCP- and METH-induced cognitive deficits. Both drugs cause the same PL neurons to acquire a new transmitter phenotype characterized by expression of GABA, GAD67, and VGAT, combined with lower levels of VGLUT1. Other PL neurons show the same transmitter phenotype in drug naïve conditions, as suggested by earlier studies^45,46^. This change in transmitter phenotype causes locomotor sensitization and memory deficits in the NORT and SAT, consistent with the involvement of the PL and the NAc, which receives input from PL neurons that change their transmitter identity^24,25,29–31,47^. Given the role of GABAergic long-range projections in modulating brain oscillation and synchronization^48^, gain of GABA by PL neurons projecting to the NAc may contribute to the reduction of NAc firing rates and disruption of cortex-accumbens synchronization after PCP-treatment ^49^.

Neuronal activity mediates this drug-induced change in transmitter identity, as expected for activity-dependent neurotransmitter switching^13,32,33^, and is necessary to maintain the newly acquired transmitter phenotype after the end of drug-treatment. Midbrain cholinergic neurons that change transmitter identity in response to sustained exercise spontaneously revert to expression of their original transmitter within a week of cessation of the stimulus^13^. In contrast, PL glutamatergic neurons maintain their GABAergic phenotype for more than 3 weeks after the end of drug-treatment and the linked cognitive deficits are long-lasting^18,37^. c-fos expression in the PL increases after acute treatment with PCP or METH^38,50^, as well as after one hour of phasic stimulation of VTA dopaminergic neurons, and remains elevated for at least two weeks^51,52^. Suppressing drug-induced hyperactivity during or after drug-treatment, respectively, prevents or rescues the switch in transmitter phenotype and the coupled behavioral alterations, indicating that PL hyperactivity is necessary to produce and maintain these changes. Because exposure to either PCP or METH decreases expression of PV and GAD67^22,53^, impaired function of PFC PV^+^ interneurons may contribute to PL glutamatergic neuron hyperactivity and maintenance of the newly acquired transmitter phenotype. Chronic treatment with clozapine also reduces c-fos expression in the PL of PCP-treated mice and reverses PCP-induced changes in transmitter phenotype and behavior. This effect may be mediated by increased inhibitory input to PL glutamatergic neurons^54^.

Signaling by dopaminergic neurons in the VTA is necessary for PL neurons to change their transmitter identity, because suppression of dopaminergic hyperactivity in the VTA during PCP- or METH-treatment prevents it. Optogenetic stimulation of phasic firing of dopaminergic neurons in the VTA is sufficient to cause PL hyperactivity and induce the switch in transmitter phenotype. Because VTA dopaminergic neurons can release transmitters other than dopamine^55^, it remains to be determined whether the change in transmitter phenotype of PL neurons is mediated by dopaminergic signaling alone or if other transmitters are also involved. Many addictive substances promote phasic firing of these dopaminergic neurons^56^. Reproducing this firing pattern optogenetically enhances DA release in the NAc^57^ and replicates some of the neuroplastic and behavioral effects of cocaine, including conditioned place preference and acquisition of self-administration that can lead to compulsive-like behavior^44,57^. This stimulation only models some aspects of drug intake^58^ and does not account for the non-dopaminergic mechanisms through which multiple drugs of abuse differentially impact brain function and behavior. Evidence that multiple drugs of abuse acutely promote mPFC hyperactivity^59,60^, and that stimulation of VTA dopaminergic neurons is sufficient to change the transmitter identity of PL neurons, raises the possibility that drugs other than METH and PCP may induce cognitive deficits by switching the transmitter phenotype of PL glutamatergic neurons.

## Methods

### Mice

All animal procedures were carried out in accordance with NIH guidelines and approved by the University of California, San Diego, Institutional Animal Care and Use Committee. Mice were maintained on a 12 h:12 h light:dark cycle (light on: 7:00 am–7:00 pm) with ad libitum access to food (7912.15 Irradiated Teklad LM Mouse/Rat diet) and water. Temperature was maintained between 65 and 75 °F (∼18 and 23 °C) with 40–60% humidity. Mice were preferentially housed 3 or 4 per cage with nesting material. After receiving surgery, mice were single-housed with nesting material. All experiments were performed on 8- to 14-week-old male mice.

C57BL/6J mice (JAX#000664) (referred to as “wild-type”), VGLUT1-IRES2-Cre mice (JAX# 037512) (referred to as “VGLUT1^CRE^”), Rosa26-LSL-H2B-mCherry (JAX#023139) (referred to as “mCherry”), B6 PV^cre^ mice (JAX#017320) (referred to as “PV^CRE^”), Ai14(RCL-tdT)-D mice (JAX#007914) (referred to as “TdTomato”), Slc32a1-2A-FlpO-D knock-in mice (JAX# 029591) (referred to as “VGAT^FLP^”), Ai65(RCFL-tdT) mice (JAX#021875) (referred to as “TdTomato^cON/fON^”), CAGGCre-ER^TM^ mice (JAX# 004682) (referred to as “CreER^T^”) were obtained from Jackson Laboratories. DAT-IRES-Cre mice (JAX#006660) (referred to as “DAT^CRE^”) were provided by the Davide Dulcis, Thomas Hnasko, and Cory Root laboratories.

Heterozygous VGLUT1^CRE^ animals were bred with either wild-type mice to obtain VGLUT1^CRE^ mice, or with homozygous mCherry mice to obtain VGLUT1^CRE^::mCherry experimental mice. PV^CRE^ mice were maintained in homozygosis and bred with either TdTomato homozygous mice to obtain PV^CRE^::TdTomato mice or wild-type animals to obtain PV^CRE^ heterozygous experimental mice.

To obtain VGAT^FLP^::CreER^T^::TdTomato^cON/fON^ mice we first bred heterozygous CreER^T^ mice with either heterozygous TdTomato^cON/fON^ or heterozygous VGAT^FLP^ animals to obtain CreER^T^::TdTomato^cON/fON^ and VGAT^FLP^::CreER^T^ breeders. These breeders were then crossed with VGAT^FLP^ or TdTomato^cON/fON^ mice, respectively, to obtain VGAT^FLP^::CreER^T^::TdTomato^cON/fON^ experimental mice.

Homozygous DAT^CRE^ mice were bred with wild-type animals to obtain DAT^CRE^ heterozygous experimental mice.

### Drugs and pharmacological treatments

Phencyclidine hydrochloride (PCP; Sigma-Aldrich, P3029) was dissolved in sterile saline and administered subcutaneously (s.c.) at a dose of 10 mg/kg/day for 10 consecutive days^13^. Methamphetamine hydrochloride (METH; Sigma-Aldrich, M8750) was dissolved in saline and administered s.c. at a dose of 1 mg/kg/day for 10 consecutive days (modified from^25^). To investigate whether drug-treatment affects the transmitter identity of PL neurons, mice were sacrificed either 2 days (Fig. 1c,g,h, Extended Data Fig. 1c-g,j,k, Fig. 2c,e,f, Extended Data Fig. 4a,b,g, Fig. 6d), 11 days (Fig. 1k, Extended Data Fig. 3c, Fig. 3d), 17 days (Fig. 4c) or 23 days (Fig. 5c) after the end of PCP or METH treatment. To investigate the effect of PCP or METH on PL neuronal activity, mice were sacrificed 2h after a single injection of the drug (Extended Data Fig. 5c-e), 2h after the last of 10 daily injections (Extended Data Fig. 5f-h) or 2 days after the end of 10-days drug-treatment (Extended Data Fig. 5i-k).

Tamoxifen (Sigma-Aldrich, T5648) was dissolved in corn oil (Sigma-Aldrich, C8267)/ethanol 9:1 and administered intraperitoneally (i.p.) at a dose of 75 mg/kg/day. To test whether PCP and METH affect the neurotransmitter phenotype of the same PL neurons, VGAT^FLP^::CreER^T^::TdTomato^cON/fON^ mice received a 10-day treatment with either saline or PCP (10 mg/kg/day). Beginning on day 9 of saline/PCP administration, mice received an injection of tamoxifen each day at 2 pm for 7 consecutive days. Mice were then left untreated for 7 days to enable tamoxifen washout before exposing them to additional treatments^61^. After the end of the washout period, mice received PCP, METH, or saline for 10 additional days and were sacrificed 2 days after the last injection.

PSEM^308^ hydrochloride (Fisher Scientific, 64-252-5) was dissolved in DMSO to obtain a stock solution of 5 mg/ml. Before use, the stock solution was further diluted in saline to a final concentration of 1 mg/ml. To investigate the effect of acute chemogenetic manipulation on neuronal activity, mice were injected i.p. with PSEM^308^ (5 mg/kg) in DMSO/saline (1:4) or DMSO/saline alone (vehicle) 10 min prior to injecting METH, PCP or saline, and were sacrificed 2h after drug-injection. To investigate the effect of repeated chemogenetic manipulation mice were injected i.p. twice a day for 10 days with either PSEM^308^ (5 mg/kg at 10-11 am and 2.5 mg/kg at either 1-2 pm or 4-5 pm) or vehicle. For experiments in which PSEM^308^ was administered together with PCP or METH, the first PSEM^308^ injection was administered 10 min prior to injecting METH, PCP or saline, and the second PSEM^308^ injection occurred 3 h later. For experiments in which the pharmacogenetic treatment occurred after the end of drug treatment, the two PSEM^308^ daily injections were administered 6 h apart.

Clozapine (Sigma-Aldrich, C6305) was dissolved in DMSO to obtain a stock solution of 10 mg/ml. The stock solution was further diluted in sterile saline before use to a final concentration of 0.5 mg/ml. DMSO dissolved 1:19 in saline (vehicle) was used as a control solution. Three days after the end of PCP treatment, mice began receiving daily i.p. injections of clozapine (5 mg/kg/day) or DMSO/saline (1:19) alone (vehicle) for 2 additional weeks^18^.

### Immunohistochemistry

Mice were deeply anesthetized with isoflurane vapor at the sacrifice time point and transcardially perfused with phosphate-buffered saline (PBS) followed by 4% paraformaldehyde (PFA) in PBS. Brains were dissected and post-fixed in 4% PFA overnight (o/n) at 4 °C, before being transferred to 30% sucrose in PBS for 2 days at 4 °C. Coronal sections, 30 µm in thickness, were obtained using a freezing microtome (Leica SM2010R) and stored at -20 °C in a cryoprotectant solution (30% glycerol, 30% ethylene glycol, 20% 0.2 M phosphate buffer).

For immunostaining, sections were washed to remove residues of cryoprotectant solution and permeabilized in 0.3% Triton X-100 in PBS. After a 2 h-incubation period in a blocking solution (5% normal horse serum, 0.3% Triton X-100 in PBS) at 22 °C, sections were incubated o/n on a rotator at 4 °C with primary antibodies diluted in the blocking solution. After washing in 0.3% Triton X-100 in PBS (three times, 15 min each), sections were incubated for 2 h on a rotator at 22 °C with secondary antibodies diluted in blocking solution. After additional washings (three times, 15 min each) in 0.3% Triton X-100 in PBS, sections were mounted with Fluoromount-G (Southern Biotech) containing DRAQ-5 (Thermo Fisher, 62251, 1:1000 dilution) when nuclear staining was needed.

Primary antibodies used in this study were:

rabbit-anti-GABA (Sigma-Aldrich, A2052, RRID:AB_477652, 1:1000), guinea pig-anti-GABA (Sigma-Aldrich, AB175, RRID:AB_91011, 1:500), mouse-anti-GAD67 (Millipore, MAB5406, RRID:AB_227872, 1:500), goat-anti-doublecortin (Santa Cruz, sc-8066, RRID:AB_2088494, 1:300) and rabbit-anti-Ki67 (Cell Signaling, 9129, RRID:AB_2687446, 1:300), rabbit-anti-RFP (Avantor VWR, RL600-401-379, RRID:AB_2209751, 1:1000), mouse-anti-cFos (Abcam, ab208942, RRID:AB_2747772, 1:500), rabbit-anti-cFos (Abcam, ab214672, 1:1000), rabbit-anti-PV (Swant, PV27, RRID:AB_2631173, 1:2000), mouse-anti-PV (Millipore, P3088, RRID:AB_477329, 1:1000), rabbit-anti-GFP (Thermo Fisher, A11122, RRID:AB_221569, 1:1000), chicken anti-GFP (Abcam, ab13970, RRID:AB_300798, 1:1000), chicken-anti-mCherry (Abcam, ab205402, RRID:AB_2722769, 1:2000), rabbit-anti-fluorescent gold (Sigma-Aldrich, RRID:AB_2632408, AB153-I, 1:500), mouse-anti-TH (Millipore, MAB318, RRID:AB_2201528, 1:1000), rabbit-anti-TH (Millipore, AB152, RRID:AB_390204, 1:2000).

Secondary antibodies for immunofluorescence were used at a concentration of 1:500. The following antibodies were from Jackson Immuno Research: Alexa Fluor-488 donkey-anti-rabbit (705-545-003, RRID:AB_2340428), Alexa Fluor-647 donkey-anti-rabbit (711-605-152, RRID:AB_2492288), Alexa Fluor-488 donkey-anti-mouse (715-545-150, RRID:AB_2340846), Alexa Fluor-488 donkey-anti-goat (705-545-147, RRID:AB_2336933), Alexa Fluor-594 donkey-anti-mouse (715-585-150, RRID:AB_2340854), Alexa Fluor-647 donkey-anti-mouse (715-605-150, RRID:AB_2340862), Alexa Fluor-488 donkey-anti-chicken (703-545-155, RRID:AB_2340375), Alexa Fluor-594 donkey anti-chicken (703-585-155, RRID:AB_2340377), Alexa Fluor-488 donkey anti-guinea pig (706-545-148, RRID:AB_2340472), Alexa Fluor-594 donkey anti-guinea pig (706-586-148, RRID:AB_2340475). The following secondary antibodies were from Life Technologies: Alexa Fluor-555 goat-anti-mouse (A21422, RRID:AB_2535844), Alexa Fluor-555 goat-anti-rabbit (A21429, RRID:AB_2535850).

### TUNEL assay

TUNEL assay was used to detect *in situ* apoptosis. The assay was performed using the In Situ Cell Death Detection (TUNEL) Kit with TMR Red (Roche, 12156792910) as previously described^13^. Briefly, 30 µm thick PL sections were fixed with 1% PFA for 20 min at 22–24 °C and rinsed with PBS (three times, 5 min each). Sections were then permeabilized in 0.1% sodium citrate and 1% Triton X-100 for 1 h at 22–24 °C. After rinsing in PBS (three times, 5 min each), sections were incubated with TUNEL reaction solution according to the vendor’s instructions. Incubation was performed in a humidified chamber for 3 h at 37 °C in the dark. Sections were rinsed and mounted with Fluoromount containing DRAQ-5 (1:1000). As positive controls, sections were treated with DNase I (10 U/mL, New England Biolabs, M0303S) for 1 h at 37 °C, rinsed in PBS (three times, 5 min each), and incubated with the TUNEL mixture.

### Fluorescent *in situ* hybridization (FISH)

FISH was performed using the RNAscope Multiplex Fluorescent v2 Kit (Advanced Cell Diagnostics, 323100) according to the manufacturer’s instructions, with a few adjustments. Briefly, the 30-μm fixed brain sections were mounted on Superfrost Plus slides and air-dried in a 50 °C oven for 30 min. Sections were rehydrated in PBS for 2 min and incubated for 5 min in 1X target retrieval solution at 95 °C. After one rinse in distilled water (2 min), sections were dehydrated with 100% ethanol for 5 seconds, air-dried, and incubated for 10 min in 5% hydrogen peroxide at RT. Sections were then incubated in a HybEZ humidified oven at 40 °C with protease III for 30 min, and later with the probe solution for 2 h. After incubation with the probes, slides were incubated o/n in a solution of SSC5X at 22 °C. The following day, sections were incubated with the following solutions in a HybEZ humidified oven at 40 °C with three rinsing steps in between each: amplification Amp-1, 30 min; Amp-2, 30 min; Amp-3, 15 min. For each probe used, sections were incubated in a HybEZ humidified oven at 40 °C with the following solutions: HRP-C1, -C2, or -C3 (depending on the probe) for 15 min, Opal dye of choice for 30 min, and HRP-blocker for 15 min. Opal 520 (Akoya Biosciences, FP1487001KT), Opal 570 (Akoya Biosciences, FP1488001KT) and/or Opal 690 (Akoya Biosciences, FP1497001KT) dyes were used for fluorescent labelling.

Depending on experimental needs, we obtained different levels of signal amplification by changing the dilution of the Opal dyes. To achieve a fully amplified signal, the Opal dyes were used at a dilution of 1:1500. This dilution was used to identify the cell-boundaries of neurons co-expressing mCherry and GAD1 (used to detect mCherry and GAD1 transcripts in: Fig. 1e-h, Extended Data Fig. 1e,f, Fig. 2e,f) and to quantify the number of VGLUT1^+^/ GAD1^+^ PL neurons (Fig. 1k, Fig. 2h,i,j,l,m,o, Fig. 3c,d, Fig. 5b,c, Fig. 6c,d,g,h). Because VGLUT1 expression is decreased but not completely lost in neurons that have changed their transmitter identity, obtaining a fully amplified VGLUT1 signal allows detection of these cells.

To obtain unsaturated, puncta-like staining, Opal dyes were diluted 1:12000. This dilution was used to quantify changes in the expression levels of VGLUT1 and VGAT (Fig. 1e-h, Extended Data Fig. 1e,f, Fig. 2e,f).

The following probes were used: mouse Probe-Mm-Slc17a7 (VGLUT1) (Advanced Cell Diagnostics, 416631), Probe-mCherry (Advanced Cell Diagnostics, 431201), Probe-Mm-Slc32a1 (VGAT) (Advanced Cell Diagnostics, 319191), Probe-Mm-GAD1 (Advanced Cell Diagnostics, 400951).

For the experiments illustrated in (Fig. 2h,i,j,l,m,o) RNAscope was followed by standard immunofluorescent staining using rabbit-anti-RFP (Avantor VWR, RL600-401-379, 1:1000) as the primary antibody, and Alexa Fluor-647 donkey-anti-rabbit (711-605-152) as the secondary antibody.

### Imaging

Images were acquired with a Leica SP5 confocal microscope with a 25x/0.95 water-immersion objective and a z resolution of 1 μm, or with Leica Stellaris 5 with a 20x/0.75 CS2 dry objective and a z resolution of 1 μm for immunohistochemistry and 0.7 μm for RNAscope. Low magnification images were acquired with Leica Stellaris 5 with a 10x/0.40 CS2 dry objective with a z resolution of 2.5 μm.

### Cell counting

All counts were performed by investigators double-blinded to the origin of each image. Either Image-J/Fiji or Imaris9 was used for cell counting (Fig.2C, Extended Data Fig. 4a,b, Extended Data Fig. 6f,h,i, Fig. 4c, Extended Data Fig. 7g,h, Extended Data Fig. 8c,d). When using Image-J/Fiji, cell counts were performed by examining all sections within the confocal stacks without maximal projection. When using Imaris9, cell counts were performed semiautomatically using the Spot detection function and the Colocalize spot plug-in, and later corrected manually. Independently of the software used for analysis, only neurons showing colocalization in at least 3 consecutive z-planes were included in the co-expression group.

PL sections were analyzed from Bregma +2.8 mm to Bregma +1.54 mm, according to the Paxinos Mouse Brain Atlas. PL boundaries were determined based on PL cytoarchitecture, as previously described^62^. Pilot experiments determined that counting 1 in 6 sections was sufficient to estimate the number glutamatergic and GABAergic cells in the PL. Consequently, to determine the number of PL mCherry^+^/GABA^+^ or GAD67^+^ neurons (Fig. 1c, Extended Data Fig. 1c,d, Fig. 2c; Extended Data Fig. 4a,b, Fig. 4c), TdTomato^+^/PV^+^ neurons (Extended Data Fig. 1j,k), as well as VGLUT1^+^/GAD1^+^ neurons (Fig. 3d), 7-to-8 sections were counted for each mouse brain. The total number of co-expressing neurons was calculated by multiplying the number of counted cells by 6. We later determined that by counting 1 in 12 sections and multiplying the number of counted cells by 12, instead of 6, the final result was not different from that obtained by counting 1 section every 6. We therefore adopted this strategy for the rest of the counts (Fig. 1k, Fig. 2l,o, Fig. 5c, Fig. 6d,h, Extended Data Fig. 10i,j).

To check for equal sampling across experimental groups, we counted the total number of PL mCherry^+^ or VGLUT1^+^ neurons and determined that their number was constant across treatment groups (Extended Data Fig. 1d, Extended Data Fig. 4b, Extended Data Fig. 10j). For TUNEL assay, as well as quantification of DCX^+^, Ki67^+^, and c-fos^+^ cells, we scored 1 in 9 PL sections, consequently counting 4-to-6 sections for each mouse brain. To quantify c-fos, GFP, and TH expression in the VTA, we collected 1 in 6 sections from Bregma -2.92 mm to Bregma -3.88 mm, according to the Paxinos Mouse Brain Atlas, thus quantifying 4-5 sections per brain.

### Quantification of VGAT and VGLUT1 mRNA expression

To quantify the expression level of VGLUT1 and VGAT mRNA, we used multiplex RNAscope against mCherry, GAD1, and either VGLUT1 or VGAT in sections from VGLUT1^CRE^::mCherry mice. The RNA signals for mCherry and GAD1 were fully amplified to allow clear detection of mCherry^+^ and/or GAD1^+^ cell boundaries. To facilitate quantification of mRNA expression levels, we obtained unsaturated, puncta-like RNA-signals for VGLUT1 and VGAT.

After staining, 4-to-7 optical sections (1 µm z step) of each physical section were examined and regions of interest (ROIs) were drawn around the boundaries of mCherry^+^, GAD1^+^ and mCherry^+^/GAD1^+^ cells using the optical section in which the cross-sectional area of the cell was the largest.

VGLUT1 or VGAT expression was quantified using Image-J/Fiji as percent of the ROI occupied by VGLUT1 or VGAT RNA fluorescent signal. For each mouse, we analyzed the ROIs of 25 cells co-expressing mCherry and GAD1, 25 cells expressing only mCherry, and 25 cells expressing only GAD1. These cells were found in sections at different levels of the PL rostrocaudal axis and were distributed across all layers of the PL.

### Stereotaxic injections

4-5-week-old mice were deeply anesthetized using 3-4% vaporized Isoflurane and head-fixed on a stereotaxic apparatus (David Kopf Instruments Model 1900) for all stereotaxic surgeries. Anesthesia was maintained throughout the procedure at a level that prevented reflex response to a tail/toe pinch, using a continuous flow of 1-2% vaporized Isoflurane. Eye drops (Puralube Vet Ointment, Fisher Scientific, 2024927) were placed in each eye to prevent them from drying out, and vitals were checked every 10 min. An incision was made to expose the bregma and lambda point of the skull. A 1 mm drill was used to perforate the skull at the desired coordinates. Stereotaxic coordinates for the injection sites were determined using the Paxinos Brain atlas and adjusted experimentally. Using a syringe pump (PHD Ultra™, Harvard apparatus, no. 70-3007) installed with a microliter syringe (Hampton, 1482452A) and capillary glass pipettes with filament (Warner Instruments, G150TF-4), we infused the brain with 500 nl (for injections in the PL) or 1 μl (for injections in the VTA) of AAV solution for each injection site at a rate of 100 nl/min. To guarantee sufficient AAV expression across the anterior-posterior (AP) extent of the PL the AVV solutions were injected bilaterally at 2 injection sites for each brain hemisphere (from bregma: anterior– posterior (AP), +2.65 mm and +2.25 mm from bregma; mediolateral (ML), ±0.5 mm; dorsal–ventral (DV), −0.8 mm and -1.1 mm from the dura).

After injection of the PL, the pipette was left in place for 8 min to allow diffusion of the virus. When the surgery was completed, the scalp was disinfected with betadine and sutured with tissue adhesive glue (Vetbond tissue adhesive, 1469SB). For post-op pain treatment, mice received an injection of Buprenorphine SR (0.5 mg/kg) or Ethiqa XR (3.25 mg/kg).

When targeting the VTA, we injected the AAV solutions bilaterally (from bregma: AP, -3.2 mm; ML, ±0.5 mm; DV, -4.0 mm from the dura). After injection of the VTA, the pipette was left in place for 16 min to allow diffusion of the virus. Mice that later received optogenetic VTA stimulation were implanted during the same surgical procedure with a fiber optic cannula with Ceramic Ferrule (RWD Life Sciences, R-FOC-F200C-39NA). We used the same coordinates used for AAV injection with the following modifications: the fiber was implanted at a 10° angle, and the DV coordinate was reduced to -3.9. To secure the implant to the skull, the skull was covered with a layer of OptiBond XTR Primer (OptiBond XTR Bottle Primer - 5 ml Bottle. Self-Etching) followed by OptiBond XTR Bottle Universal Adhesive (OptiBond XTR Bottle Universal Adhesive 5 ml Bottle. Self-Etching, Light-Cure). Finally, a thick layer (up to 0.5 cm thick) of Nano-optimized Flowable Composite (Tetric EvoFlow A2 Syringe - Nano-optimized Flowable Composite 1 - 2 Gram) was used to create a scaffold and secure the optic fiber to the skull. Polymerization of OptiBond XTR Primer, Adhesive, and Flowable Composite was achieved with dental LED light (Fencia Premium Silver LED Light, 5W).

### Viral constructs

To suppress GAD67 expression in PL glutamatergic neurons we used AAV9-CAG-DIO-shRNAmir-scramble-GFP (6.30E+13 particles/ml), and AAV9-CAG-DIO-shRNAmir-mGAD1-GFP as control (2.41E+14 viral particles/ml). pAAV-CAG-DIO-shRNAmir-mGAD1-EGFP and pAAV-CAG-DIO-shRNAmir-Scramble-EGFP plasmid ^13^ were produced by Vector Biolabs and AAV9 vectors were packaged in the Salk Institute Viral Vector Core. The shRNA sequence for mouse GAD1 is 5′-GTCTACAGTCAACCAGGATCTGGTTTTGGCCACTGACTGACCAGATCCTTTGACTGTAGA-3′.

Validation of AAV-CAG-DIO-shRNAmir-scramble-GFP and AAV-CAG-DIO-shRNAmir-mGAD1-GFP can be found in^13^.

To activate PL PV^+^ neurons we used AAV9-syn-FLEX-rev-PSAML141F,Y115F:5HT3HC-IRES-GFP (4.61E+12 viral particles/ml). rAAV-syn::FLEX-rev::PSAML141F,Y115F:5HT3HC-IRES-GFP was a gift from Scott Sternson (Addgene plasmid # 32477; http://n2t.net/addgene:32477; RRID:Addgene_32477)^63^, and the AAV9 vector was packaged in the Salk Viral Vector Core.

To inhibit dopaminergic neurons in the VTA, we used AAV9-syn::FLEX-rev::PSAML141F,Y115F:GlyR-IRES-GFP (2.1E+12 viral particles/ml). rAAV-syn::FLEX-rev::PSAML141F,Y115F:GlyR-IRES-GFP was a gift from Scott Sternson (Addgene plasmid # 32481; http://n2t.net/addgene:32481; RRID:Addgene_32481)^63^.

To optogenetically stimulate VTA dopaminergic neurons, we used AAV5-EF1a-double floxed-hChR2(H134R)-EYFP-WPRE-HGHpA (1E+12 viral particles/ml) and AAV5-Ef1a-DIO EYFP as control (2.3E+12 viral particles/ml). pAAV-EF1a-double floxed-hChR2(H134R)-EYFP-WPRE-HGHpA and pAAV-Ef1a-DIO EYFP were a gift from Karl Deisseroth (Addgene viral prep # 20298-AAV5; http://n2t.net/addgene:20298; RRID:Addgene_20298; and Addgene viral prep # 27056-AAV5; http://n2t.net/addgene:27056 ; RRID:Addgene_27056).

### Retrograde tracing

For retrograde tracing, 50 nl of Fluoro-gold (Hydroxystilbamidine Fluoro-Gold™), 4% in H_2_O, (Biotium, 80023) were stereotaxically injected at 50 nl/min into the ventral nucleus accumbens (from bregma: AP, +1.40 mm; ML, ±1.20 mm; and DV, −4.20 mm from the dura). The glass pipette was left in place for 10 min after injection.

### Optogenetic stimulation

The first stimulation session occurred 4-5 weeks after surgery. After being moved to the room where stimulation was performed, mice were acclimatized to the room for 1 h. Stimulation occurred in the home cage while the mouse was allowed to move. The ceramic ferrule protruding from the animal’s head was coupled to a DPSS blue light laser (473 nm Blue DPSS Laser with Fiber Coupled, BL473T3-050FC, with ADR-700A Power Supply; Shanghai Laser & Optics Century) via custom-made patch cords. Patch cords were assembled by epoxying optical fibers (200 μm, 0.39 numerical aperture, Thorlabs, FT200EMT) to Fiber Connectors FC/PC with Ceramic Ferrule (Thorlabs, 30140E1) and polishing the optic fiber with a fiber polishing kit (Thorlabs) to achieve a minimum of 85% transmission. The laser power was measured before each mouse/stimulation session using a Compact Power and Energy Meter Console, Digital 4” LCD (Thorlabs, PM 100D). Before each mouse/stimulation session, we attached an unused fiber optic cannula to the optic cable and measured the laser power at the tip of the fiber optic cannula using the Compact Power and Energy Meter Console, Digital 4” LCD and adjusted the laser power to 8.5 ± 1 mW at the tip of the fiber optic cannula. Each session of optogenetic stimulation lasted 1h, during which 80 sets of laser stimulation, each in turn consisting of 30 bursts of 5 pulses of 4 ms at 20 Hz, were delivered (modified from ^44^). For c-fos quantification experiments, mice received a single session of optogenetic stimulation and were sacrificed 1 h after the end of the session. For the 10-days stimulation protocol, mice received a daily session of 1 h stimulation between 11 am and 5pm and were sacrificed 2 days after the last stimulation.

### Drug-induced locomotor sensitization

Locomotor activity was measured as total distance traveled in the home cage during the 90 min immediately after a single injection of PCP, METH or saline. Mouse movements were recorded using a camera suspended 2 m above the home cage and were automatically scored using AnyMaze 5.2 (Stoelting, Wood Dale, IL, USA). In experiments aimed at preventing drug-induced changes in neurotransmitter phenotype by combining PCP, METH, or saline treatment with chemogenetic activation of PV+ interneurons, or by suppressing the gain of GABA with shGAD1 interference, locomotor activity was measured on the first day of treatment (i.e. immediately after the first drug/saline injection, DAY1) and on the last day of treatment (i.e. immediately after the tenth drug/saline injection, DAY10).

For experiments in which mice received either clozapine treatment or chemogenetic manipulation of PL activity after the end of PCP-, METH- or saline-treatment, locomotor quantification was performed at the end of the experimental timeline, two days after the spontaneous alternation task (SAT). In this case, baseline locomotor activity was first recorded in the home cage for 90 min, after which all mice received an acute injection of PCP 10 mg/kg (*PCP challenge*) or METH 1 mg/kg (*METH challenge*).

### Novel object recognition test (NORT)

The novel object recognition test was used to assess recognition memory performance and performed following the procedure described in ^64^ with the modifications outlined below. The test was conducted in an open field box made of dark gray plastic 40 x 40 x 21 cm, dimly illuminated (30–40 lux). In the 3 days before the test, mice were acclimatized to the open field box for 5 min/day. 24 h after the last acclimatization session, mice were placed back in the open field with two identical objects and allowed to explore the 2 objects for 10 minutes (familiarization phase). 24 h after the familiarization phase, memory retention was tested by placing the mouse back in the open field where one of the familiar objects had been replaced by a novel object (test phase). The mouse was left to explore the 2 objects for 12 minutes and video recordings were collected with a camera (Sony HDR-CX405) suspended 1 m above the apparatus. Because mice (including saline-treated controls) often failed to reach an accepted exploration criterion^64^ (20 s of total exploration in the 10 min of the test), we extended the time available for exploration from 10 to 12 min, and mice that did not reach the criterion in less than 12 min were excluded. Mice were also excluded if they moved or overturned one of the two objects. The time spent exploring each of the two objects (novel and familiar) was manually scored using BORIS software (version v. 5.0.1^65^) by investigators blinded to the mouse’s previous treatment-history. The mouse was considered to be exploring the object when it had its nose directed toward the object at a distance of <1 cm. Chewing or climbing on the object was not considered an exploratory behavior. Exploratory behavior was scored until the mouse reached the criterion of 20 s spent exploring the two objects. We calculated a recognition index (RI) as the percent of time spent exploring the novel object relative to the total time spent exploring both objects (novel and familiar) [RI=time exploring novel object/(time exploring novel object+ time exploring familiar object)]. Total exploration time and time to reach the criterion were also recorded.

### Spontaneous alternation task (SAT)

The spontaneous alternation task was performed according to the procedure described by^23,66^ and used to measure immediate working memory performance. The test was conducted in a T-shaped, dark gray plastic maze, composed of 3 arms 30 cm long, 9 cm wide and 20 cm tall. Mice naïve to the maze were placed at the end of one of the 3 arms, and left free to explore the maze for 8 min. Exploration was recorded with a camera suspended 1 m above the apparatus, and the series of arm entries was scored using BORIS software by investigators blinded to the mouse’s previous treatment history. The mouse was considered to have entered an arm of the maze only when both the forelimbs and hindlimbs were completely within the arm. Alternation was defined as successive entries into the three arms of the maze on overlapping triplet sets. Alternation percent was calculated as ratio of actual alternation to total possible alternations (defined as the total number of arm entries minus two), multiplied by 100. Total arm entries were also scored to compare the degree of exploration across treatment groups.

### Statistics

Statistical analyses of the data were performed using Prism 9 software. Excel Real Statistics package was used for non-parametric Aligned Rank Transform (ART) ANOVA. The Shapiro-Wilk test was used to assess whether the data were normally distributed. All statistical tests were two-tailed. Details about the number of animals and statistical tests used for each experiment are reported in the figure legends, the Extended Data figure legends and in supplementary table 1. Measurements were always taken from distinct samples, except for measurements of locomotor activity on first and last day of treatment, which were on the same mice (Fig. 1l; Extended Data Fig. 3d; Fig. 4g; Extended Data Fig. 7d; Fig. 5f,i; Extended Data Fig. 8f,i). Means and SEMs are reported for all experiments.

## Acknowledgements

We thank Dr. Cory Root, Dr. David Dulcis, and Dr. Thomas Hnasko for providing DAT-IRES-Cre mice (JAX#006660). We are grateful to Dr. Cory Root, Donghyung Lee and James Howe VI for providing the materials and guidance for the optogenetic experiments. We thank Dr. Scott Sternson for donating the rAAV-syn::FLEX-rev::PSAML141F,Y115F:5HT3HC-IRES-GFP and the rAAV-syn::FLEX-rev:: PSAML141F,Y115F:GlyR-IRES-GFP plasmid to Addgene. We thank Dr. Karl Deisseroth for donating the pAAV-EF1a-double floxed-hChR2(H134R)-EYFP-WPRE-HGHpA and pAAV-Ef1a-DIO EYFP plasmids to Addgene. We thank Alexander Glavis-Bloom for assisting with genotyping and for technical support; Hannah Kim, Ramiz Ahmed, and Tianna Hung for assisting with experimental procedures. We thank Dr. Cory Root, Dr. Davide Dulcis and Dr. Samuel Barnes for their thoughtful reviews and comments on an earlier version of this manuscript.

## Contributions

M.P. and N.C.S. conceived the study, designed the experiments, interpreted the results and wrote the manuscript. M.P. and A.H. performed the experiments and the analyses. A.T. performed and analyzed the clozapine experiments. H.-q.L and S.G. contributed to experimental design and data interpretation.

## Corresponding authors

Correspondence to Nicholas Spitzer or Marta Pratelli

