## Supplementary Materials for "Drug-induced change in transmitter identity is a shared mechanism generating cognitive deficits"

### Extended data

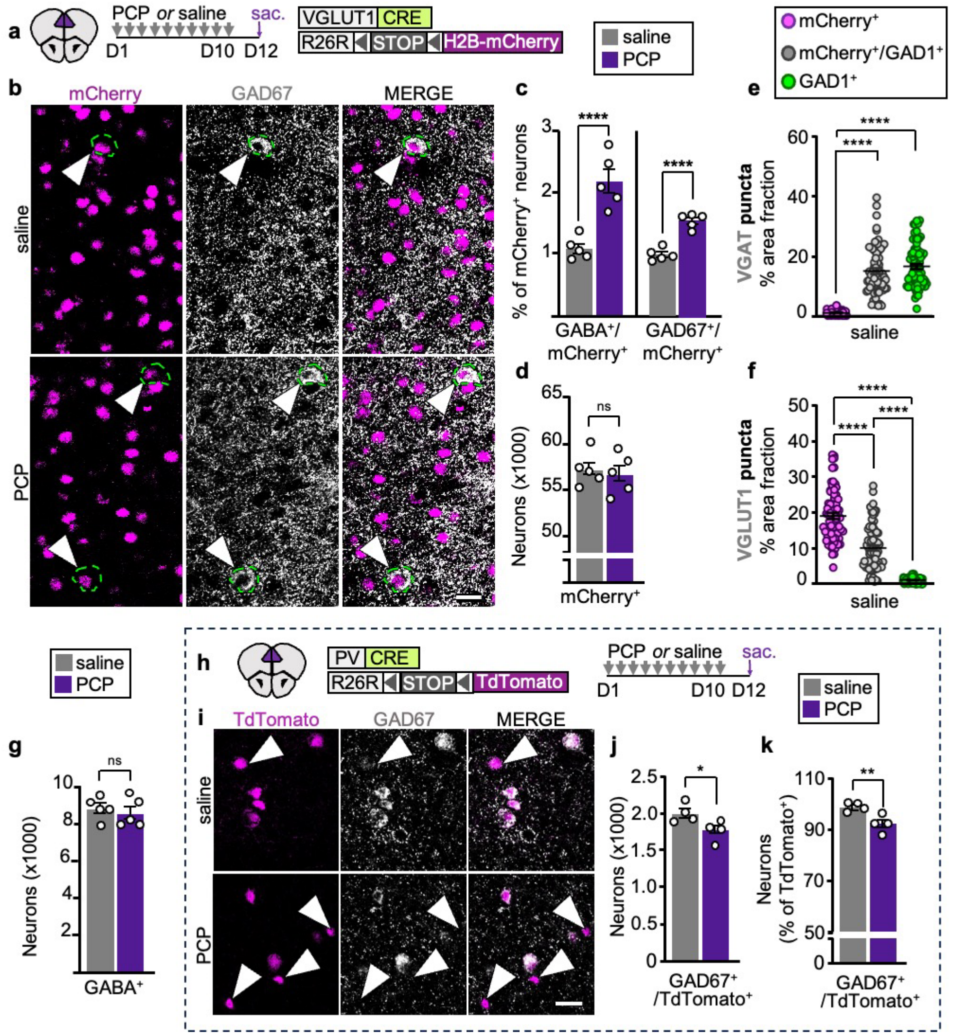

**Extended Data Fig. 1 | Expression of neurotransmitter markers in the PL of PCP-treated mice and saline controls.** (**a**) Experimental protocol to investigate the effect of PCP treatment on the transmitter phenotype of PL neurons. (**b**) PL neurons co-expressing mCherry and GAD67 (arrowheads). Scale bar, 20 µm. (**c**) Percent of mCherry$^+$/GABA$^+$ and mCherry$^+$/GAD$^+$ cells in total mCherry$^+$ neurons (*n*=5 mice). (**d**) Quantification of mCherry$^+$ neurons in the PL of PCP- and saline-treated mice (*n*=5 mice). (**e**,**f**) Quantification of expression level of VGAT (**e**) and VGLUT1 (**f**) in mCherry$^+$-only, mCherry$^+$/GAD1$^+$ and GAD1$^+$-only neurons in PL neurons of saline-treated controls (n=25 cells/type/mouse for 3 mice). (**g**) Quantification of PL neurons expressing GABA but not mCherry (*n*=5 mice). (**h**) Experimental protocol to investigate the effect of PCP treatment on GAD67 expression in PV$^+$ neurons. (**i**) PL TdTomato$^+$ labeling of PV$^+$ interneurons, and GAD67$^+$ cells across treatment groups. Arrowheads show TdTomato$^+$ neurons lacking GAD67 expression. Scale bar, 30 µm (**j**). Quantification of TdTomato$^+$/GAD67$^+$ neurons in the PL of PCP- and saline-treated mice (*n*=4 mice). (**k**) Percent of TdTomato$^+$/GAD67$^+$ cells in total TdTomato$^+$ neurons (*n*=4 mice). Statistical significance (*P<0.05, **P<0.01, ****P<0.0001) was assessed using unpaired t-test (**c**,**d**,**g**,**j**,**k**) and Kruskal-Wallis followed by Dunn's test (**e**,**f**). Data are presented as mean ± SEM. See Supplementary Table 1 for detailed statistics.

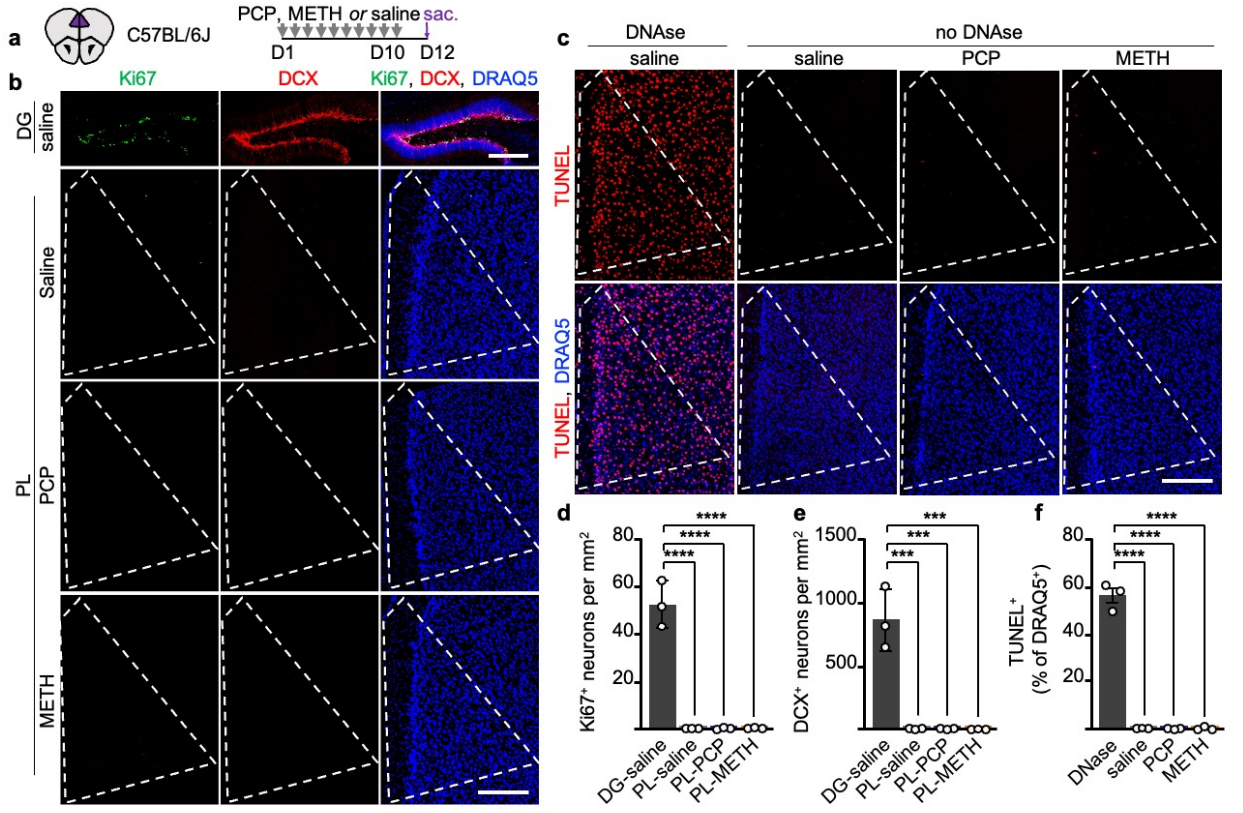


**Extended Data Fig. 2 | Neurogenesis and apoptosis are not detected in the PL of PCP- or METH-treated mice.** (**a**) Experimental protocol to evaluate whether drug-treatment causes apoptosis or neurogenesis in the PL. (**b**) Ki67, DCX, and DRAQ5 (nuclear staining) expression in the dentate gyrus (DG; upper panels) and PL (lower panels) of mice treated for 10 days with PCP, METH, or saline. Scale bar, 300 µm. (**c**) PL sections from PCP-, METH- or saline-treated mice double-stained for TUNEL and DRAQ5. Left panels contain a DNase-treated control section. Scale bar, 300 µm. (**d**,**e**) Quantification of $Ki67^+$ and $DCX^+$ cells in the DG and PL (*n*=3 mice). (**f**) Quantification of TUNEL signal in the PL of mice treated with PCP, METH, or saline (*n*=3 mice). Statistical significance (***P 0.001; ****P<0.0001) was assessed using one-way ANOVA with Tukey's multiple-comparisons test. Data are presented as mean ± SEM. See Supplementary Table 1 for detailed statistics.

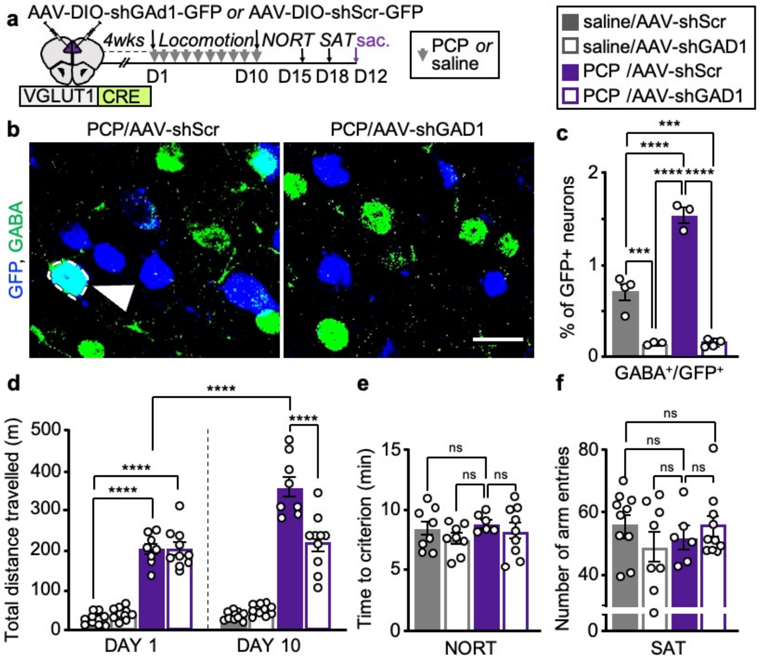


**Extended Data Fig. 3 | Behavioral effect of shGAD1 suppression of PCP-induced gain of GABA in PL glutamatergic neurons.** (**a**) Experimental protocol to test whether overriding PCP-induced gain of GABA interferes with PCP-induced behavioral changes. (**b**) GABA is co-expressed in shScr$^+$ but not shGAD1$^+$ neurons (identified by GFP expression) in the PL. Scale bar, 20 µm. (**c**) shGAD1 suppresses GABA expression in glutamatergic neurons (*n*=3-4 mice). (**d**) PCP-induced locomotor sensitization is suppressed by shGAD1 (*n*=8-10 mice). (**e**,**f**) Time to criterion on the NORT and number of arm entries on the SAT are not altered by shGAD1 expression across experimental conditions (*n*=6-10 mice). Statistical significance (***P<0.001, ****P<0.0001) was assessed using two-way ANOVA with Tukey's multiple-comparisons test (**c**-**e**), and ART factorial ANOVA followed by Mann Whitney test and Bonferroni correction (**f**). Data are presented as mean ± SEM. See Supplementary Table 1 for detailed statistics.

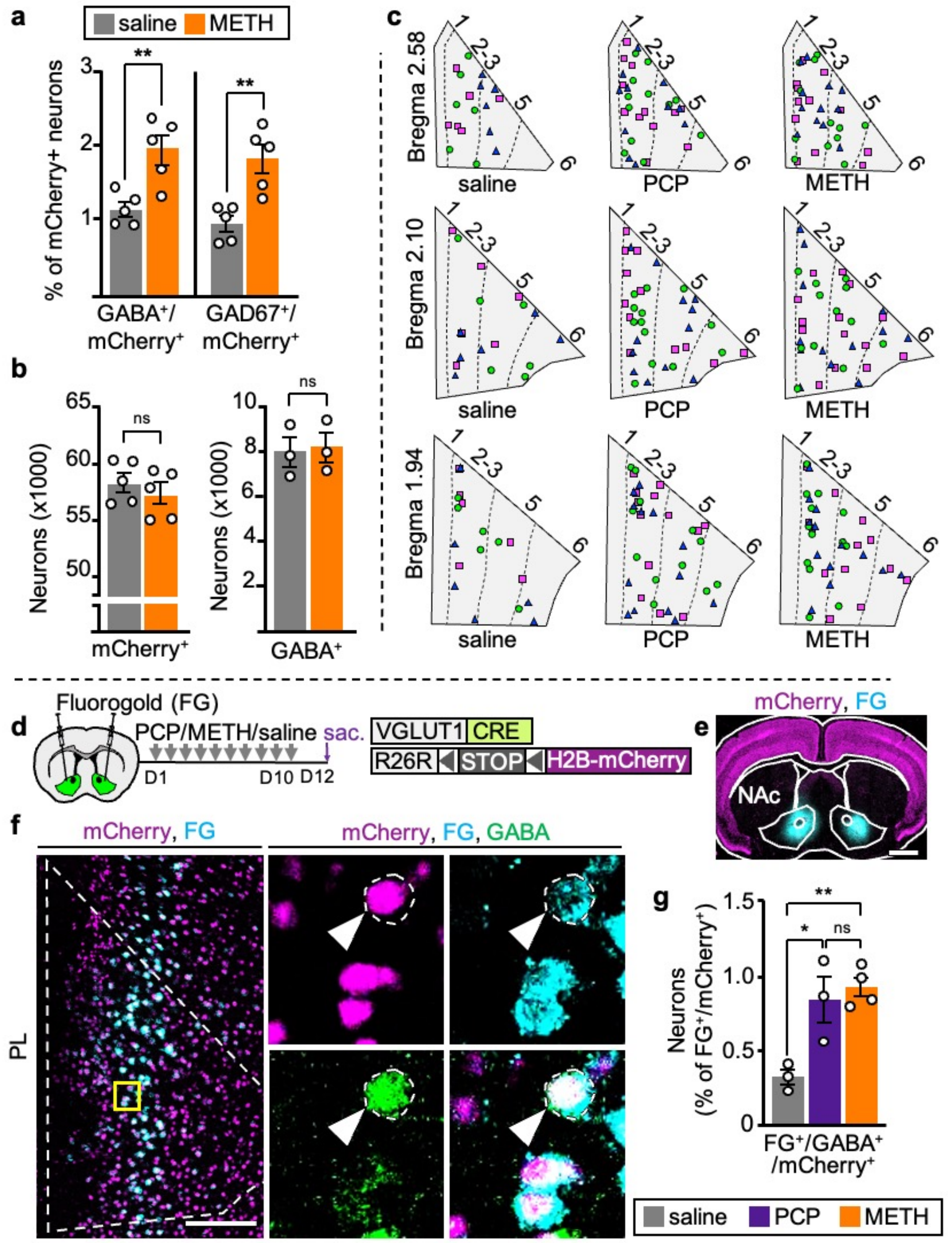


**Extended Data Fig. 4 | VGLUT1$^+$/GAD1$^+$ neurons are enriched in layer 2/3 and layer 5 of the PL and neurons that change transmitter phenotype upon drug exposure project to the nucleus accumbens.** (**a**) Percent of mCherry$^+$/GABA$^+$ and mCherry$^+$/GAD$^+$ cells in total mCherry$^+$ neurons in the PL of METH-treated mice and saline controls (*n*=5 mice). (**b**) Quantification of mCherry$^+$ and GABA$^+$/mCherry$^-$ neurons in the PL of METH-treated mice and saline controls (*n*=3-5 mice). (**c**) Cartoons show the locations across PL layers and along the PL rostro-caudal axis of glutamatergic neurons that co-express GABA without drug treatment or switch transmitter identity upon treatment with PCP or METH. Each cartoon was obtained by superimposing the location of neurons from three distinct 30 μm PL sections, each belonging to a different mouse, and all positioned at the same distance from bregma. In each cartoon, dots of the same color and shape represent cells detected in the same mouse. Cartoons in the bottom row (Bregma 1.94) are also shown in Fig. **1d**, and Fig. **2d** . L1, layer 1; L2/3, layer 2/3; L5, layer 5; L6, layer 6. (**d**) Experimental protocol to test whether PL neurons that gain GABA project to the NAc. (**e**) Fluorogold (FG) injection site in the NAc. Scale bars, 1 mm. (**f**) FG labeling in the PL. (Yellow rectangle) region illustrated at higher magnification on the right, showing a neuron co-expressing mCherry, GABA and FG. Scale bar, 200 μm. (**g**) Quantification of FG-expressing mCherry$^+$/GABA$^+$ neurons across treatments (*n*=3-4 mice). Statistical significance

(*P<0.05, **P<0.01) was assessed using unpaired t-test (**a**, **b**) and one-way ANOVA followed by Tukey's test (**g**). Data are presented as mean ± SEM. See Supplementary Table 1 for detailed statistics.

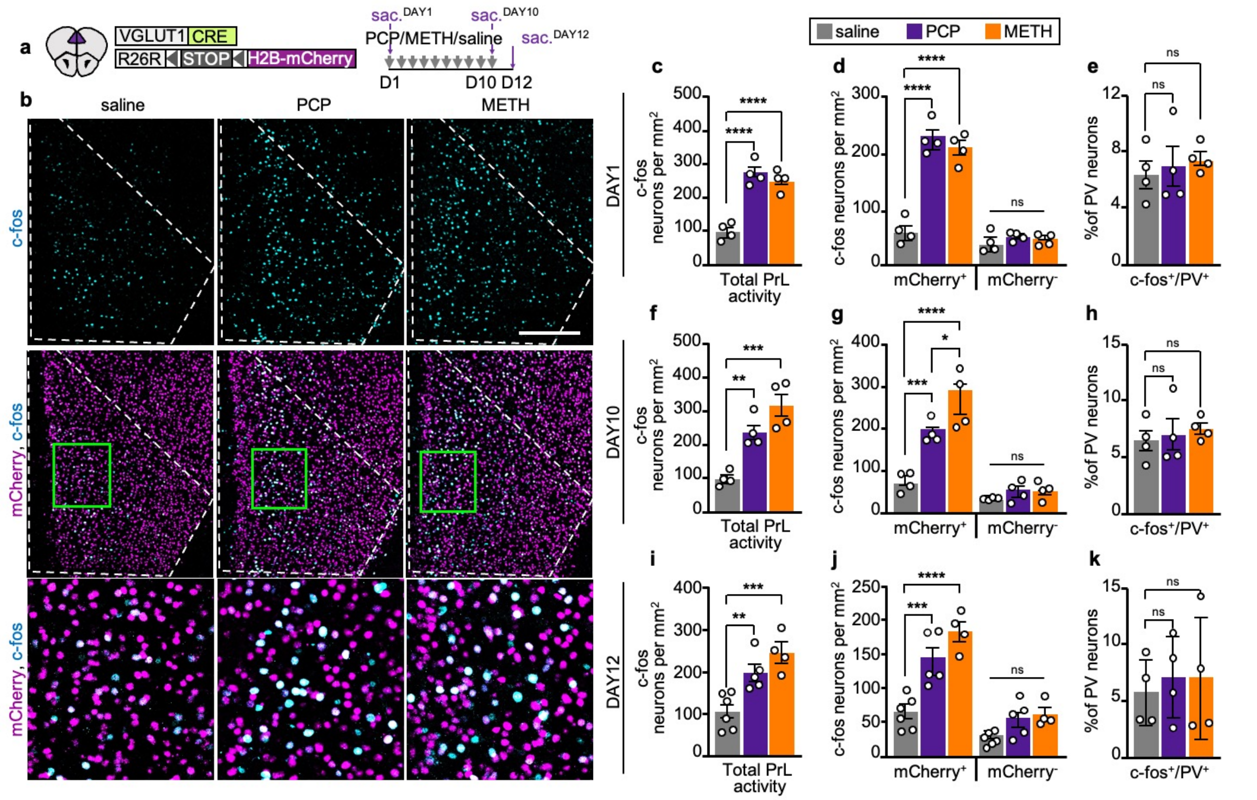


**Extended Data Fig. 5 | Both PCP and METH increase neuronal activity in PL glutamatergic neurons.** (**a**) Experimental protocol to determine the effect of drug treatment on c-fos expression in PL glutamatergic neurons. (**b**) PL c-fos-labeled mCherry$^+$ cells 2h after single drug injection on day 1. (Green rectangles) regions illustrated at higher magnification below. Scale bar, 250 μm. (**c**,**f**,**i**) Quantification of c-fos$^+$ PL neurons on day 1 (**c**) (*n*=4 mice), day 10 (**f**) (*n*=4-5 mice) and day 12 (2 days after the end of treatment) (**i**) (*n*=4-6 mice). (**d**,**g**,**j**) Quantification of c-fos labeling in mCherry$^+$ and mCherry$^-$ neurons at the same time points (*n*=4-6 mice) (**e**,**h**,**k**) c-fos labeling of PV$^+$ neurons on day 1, day 10 and day 12 (*n*=4-6 mice). Statistical significance (*P<0.05, **P<0.01, ***P<0.001, ****P<0.0001) was assessed using one-way ANOVA with Dunnett's multiple-comparisons test (**c**,**e**,**f**,**h**,**i**,**k**) or two-way ANOVA with Tukey's multiple-comparisons test (**d**,**g**,**j**). Data are presented as mean ± SEM. See Supplementary Table 1 for detailed statistics.

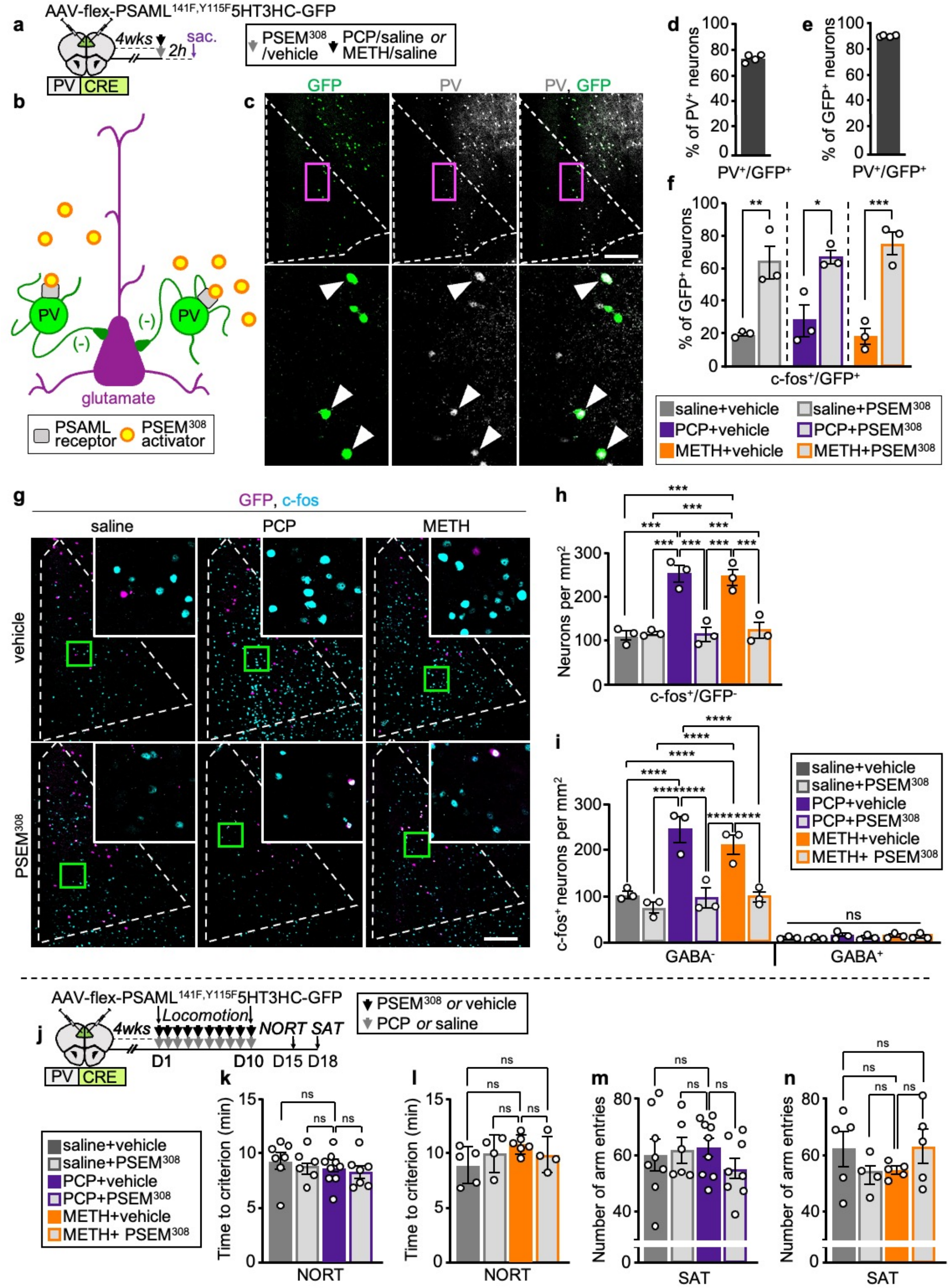


**Extended Data Fig. 6 | Chemogenetic stimulation of PV⁺ interneurons suppresses drug-induced PL hyperactivity and does not affect exploratory behaviors.** (**a**) Experimental protocol to validate AAV-flex-PSAML-5HT3HC-GFP as a chemogenetic tool to activate PL PV⁺ neurons and suppress drug-induced PL hyperactivity. (**b**) Strategy to suppress drug-induced hyperactivity of PL glutamatergic neurons. (**c**) Expression of PSAML-5HT3HC-GFP in PL PV⁺ neurons in a mouse sacrificed 2h after acute injection of METH and vehicle. (Magenta rectangles) regions shown at higher magnification below. Arrowheads indicate PV⁺ neurons expressing GFP. Scale bar, 200 μm. (**d**) Efficiency of AAV transduction measured as the percent of PV⁺/GFP⁺ neurons in the PV⁺ population (74%, 906/1229) (*n*=4 mice). (**e**) Specificity of AAV transduction measured as the percent of $PV^+/GFP^+$ neurons in the $GFP^+$ population (91%, 906/998) (*n*=4 mice). (**f**) Quantification of c-fos expression in AAV-transduced neurons (identified by GFP) 2h after injections (*n*=3 mice). (**g**) PL neurons expressing c-fos, 2h after drug injections. (Green rectangles) regions illustrated at higher magnification in insets. Scale bar, 250 μm. (**h**) Quantification of c-fos expression in PL neurons not expressing GFP (*n*=3 mice). (**i**) Quantification of c-$fos^+/GABA^+$ and c-$fos^+/GABA^-$ (ostensibly glutamatergic) PL neurons 2h after injections (*n*=3 mice). (**j**) Experimental protocol to assess the effect of chemogenetic activation of $PV^+$ neurons on exploratory behaviors. (**k-n**) Time to criterion on the NORT and number of arm entries on the SAT are not altered by chemogenetic activation of $PV^+$ neurons across experimental conditions (n=4-9 mice/group). Statistical significance (*P<0.05, **P<0.01, ***P<0.001, ****P<0.0001) was assessed using two-way ANOVA with Tukey's multiple-comparisons test (**f**,**h**,**i**,**k**-**n**). Data are presented as mean ± SEM. See Supplementary Table 1 for detailed statistics.

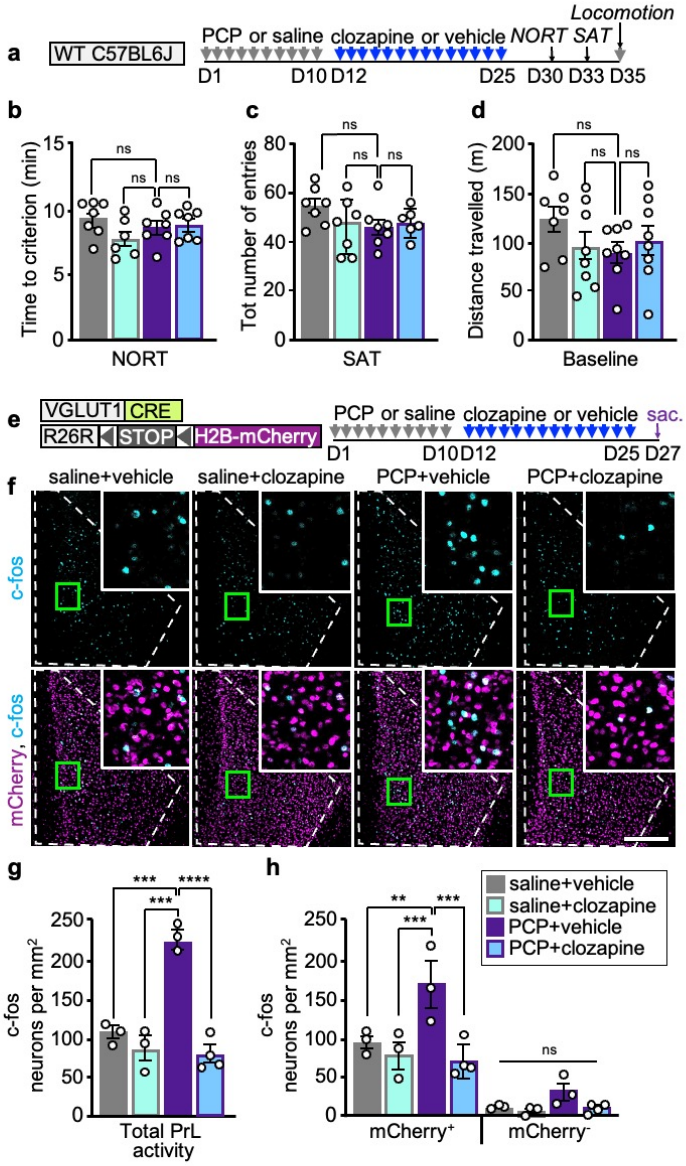


**Extended Data Fig. 7 | Clozapine treatment does not affect exploratory behaviors and normalizes PL hyperactivity after PCP-treatment.** (**a**) Experimental timeline to test the of consecutive treatment with PCP and clozapine on exploratory behaviors. (**b**-**d**) Time to criterion on the NORT, number of arm entries on the SAT and baseline locomotion are unchanged across treatment groups (*n*=6-8 mice). (**e**) Experimental protocol to determine if consecutive treatment with PCP and clozapine affects c-fos expression in the PL. (**f**) PL c-fos$^+$ neurons across treatments. (Green rectangles) regions illustrated at higher magnification in insets. Scale bar, 250 μm. (**g**,**h**) Clozapine normalizes PL c-fos$^+$ expression after the end PCP treatment (*n*=3-4 mice). Statistical significance (**P<0.01, ***P<0.001, ****P<0.0001) was assessed by two-way ANOVA with Tukey's multiple-comparisons test (**b**,**c**,**d**,**g**,**h**). Data are presented as mean ± SEM. See Supplementary Table 1 for detailed statistics.

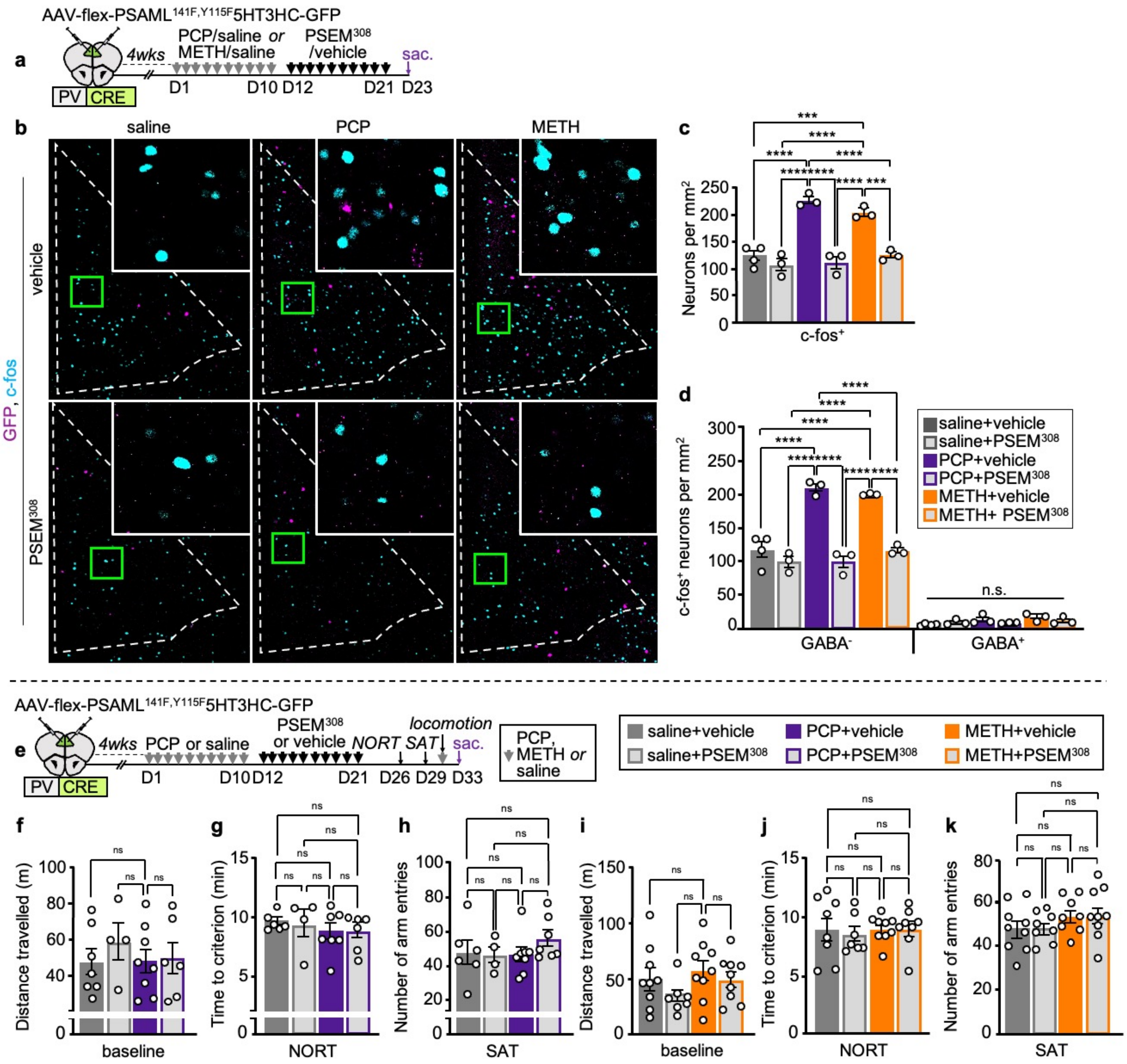


**Extended Data Fig. 8 | Chemogenetic activation of PL PV$^{+}$ neurons after the end of PCP- and METH-treatment normalizes PL activity, and does not affect exploratory behaviors.** (**a**) Experimental protocol to manipulate PL activity after the end of drug-treatment. (**b**) c-fos$^{+}$ and GFP$^{+}$ PL neurons across treatments. (Green rectangles) regions illustrated at higher magnification in insets. Scale bar, 250 μm. (**c**) c-fos expression in the PL of mice that received drug-treatment followed by 10 daily injections with PSEM$^{308}$ or vehicle as control (*n*=3-4 mice). (**d**) c-fos$^{+}$ neurons in GABA$^{+}$ and GABA$^{-}$ neurons (*n*=3-4 mice). (**e**) Experimental protocol to test the behavioral effect of repeatedly activating PL PV$^{+}$ neurons after the end of PCP-treatment. (**f**-**k**) No changes in baseline locomotion, time to criterion on NORT or number of arm entries on SAT were observed across conditions (*n*=4-9 mice). Statistical significance (***P<0.001, ****P<0.0001) was assessed using two-way ANOVA with Tukey's multiple-comparisons test (**c**,**d**,**f**-**k**). Data are presented as mean ± SEM. See Supplementary Table 1 for detailed statistics.

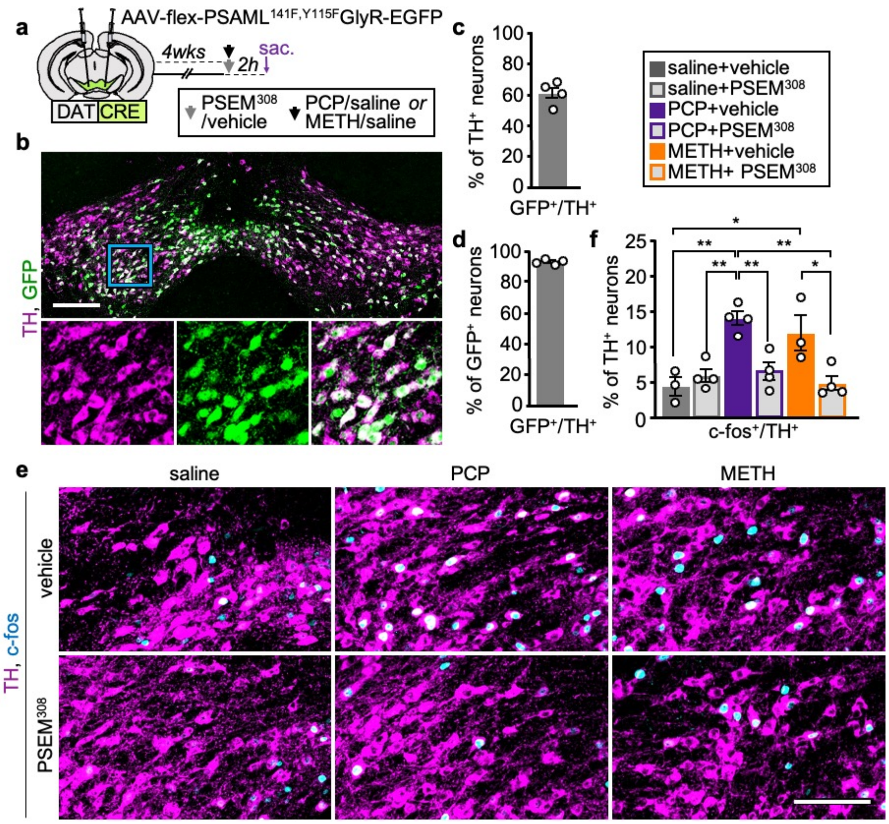


**Extended Data Fig. 9 | Chemogenetic inhibition of VTA dopaminergic neurons suppresses their activity.** (**a**) Experimental protocol to validate AAV-flex-PSAML-GlyR-GFP as a chemogenetic tool to inhibit VTA dopaminergic neurons. (**b**) Expression of TH and GFP in the VTA. (Blue rectangle) region shown at higher magnification below. Scale bar, 200 μm. (**c**) Efficiency of AAV transduction quantified as the percent of TH$^+$/GFP$^+$ neurons in the TH$^+$ population (62%, 3973/6455) (*n*=4 mice). (**d**) Specificity of AAV transduction quantified as the percent of TH$^+$/GFP$^+$ neurons in the GFP$^+$ population (92%, 3973/4297) (*n*=4 mice). (**e**) VTA expression of c-fos and TH across treatment groups. Scale bar, 200 μm. (**f**) Quantification of c-fos$^+$ expression in TH$^+$ neurons (*n*=3-4 mice). Statistical significance (*P<0.05, **P<0.01) was assessed using two-way ANOVA with Tukey's multiple-comparisons test (**f**). Data are presented as mean ± SEM. See Supplementary Table 1 for detailed statistics.

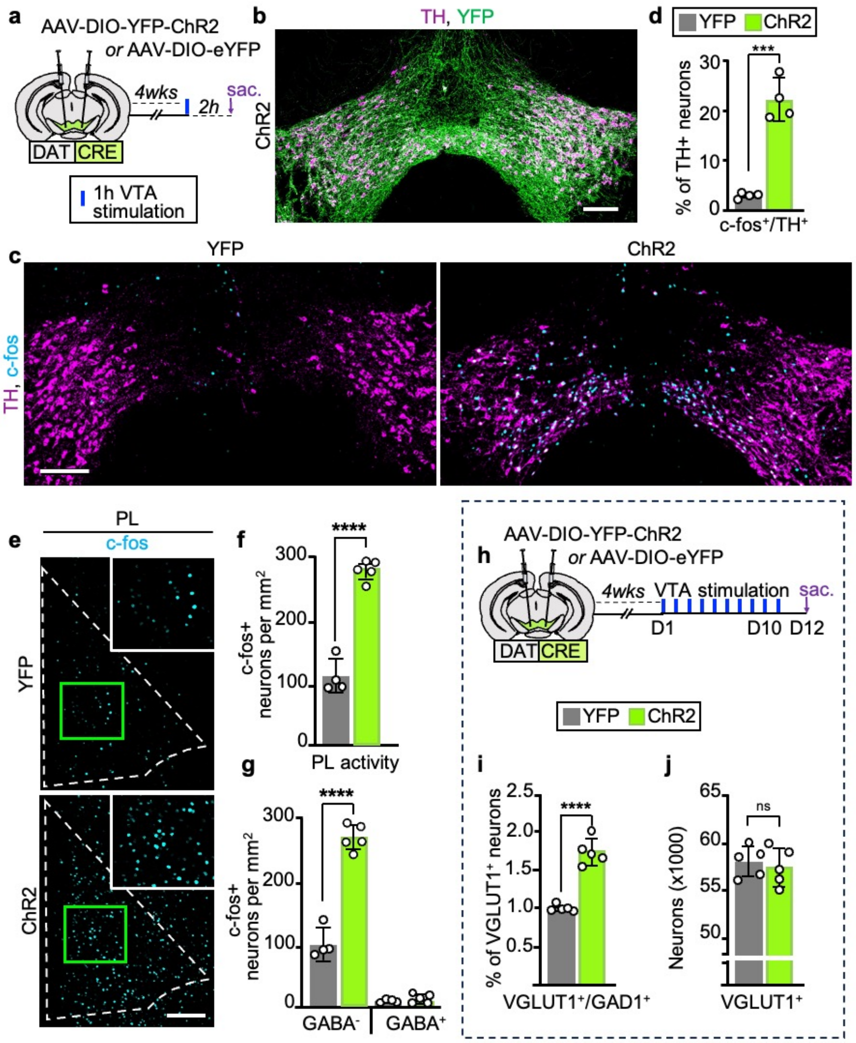


**Extended Data Fig. 10 | Optogenetic stimulation activates VTA dopaminergic neurons, increases c-fos expression in the PL and does not affect the number of PL VGLUT1⁺ neurons.** (**a**) Experimental protocol for optogenetic stimulation of VTA dopaminergic neurons. (**b**) Expression of YFP-ChR2 in the VTA. Scale bar, 200 μm. (**c**) c-fos and TH expression in the VTA following optogenetic stimulation. Scale bar, 200 μm. (**d**) Quantification of c-fos$^+$ expression in VTA TH$^+$ neurons 2h after the beginning of optogenetic stimulation (*n*=4 mice). (**e**) PL c-fos expression 2h after the beginning of VTA stimulation. (Green rectangles) regions illustrated at higher magnification in insets. Scale bar, 250 μm. (**f**) Quantification of c-fos expression in (**e**) (*n*=4-5 mice). (**g**) c-fos labeling in GABA$^+$ and GABA$^-$ neurons (*n*=4-5 mice). (**h**) Experimental protocol to investigate the effect of repeated VTA stimulation on PL neuron transmitter phenotype. (**i**) Percent of VGLUT1$^+$/GAD1$^+$ cells in total VGLUT1$^+$ neurons (*n*=5 mice). (**j**) Quantification of VGLUT1$^+$ neurons in the PL of ChR2 and YFP mice (*n*=5 mice). Statistical significance (***P<0.001, ****P<0.0001) was assessed using unpaired t-test (**d**,**f**,**i**,**j**), or two-way ANOVA with Tukey's multiple-comparisons test (**g**). Data are presented as mean ± SEM. See Supplementary Table 1 for detailed statistics.
